# A Pluripotent Developmental State Confers a Low Fidelity of Chromosome Segregation

**DOI:** 10.1101/2022.03.01.482524

**Authors:** Chenhui Deng, Amanda Ya, Duane A. Compton, Kristina M. Godek

**Author notes:** **Correspondence:** Kristina M. Godek, Ph.D., Geisel School of Medicine at Dartmouth, Department of Biochemistry and Cell Biology, HB7650, Lebanon, NH, 03756, USA.

## Abstract

Human pluripotent stem cells (hPSCs) frequently become aneuploid with abnormal chromosome numbers due to mitotic chromosome segregation errors during propagation in culture. Yet, we do not understand why hPSCs exhibit a low mitotic fidelity. Here we investigate the mechanisms responsible for mitotic errors in hPSCs and show that the primary cause is lagging chromosomes with improper merotelic chromosome microtubule attachments in anaphase. Accordingly, we can improve merotelic error correction and reduce lagging chromosome rates in hPSCs using small molecules that prolong mitotic duration or destabilize chromosome microtubule attachments providing chemical strategies to preserve genome stability. Strikingly, we also demonstrate that mitotic error rates correlate with developmental potential decreasing upon differentiation and loss of pluripotency and conversely increasing after reprogramming to a pluripotent state. Thus, chromosome segregation fidelity is inherently low in hPSCs and depends on developmental state in normal human cells.

## Introduction

Human pluripotent stem cells (hPSCs), including human embryonic stem cells (hESCs) and induced pluripotent stem cells (iPSCs), have the ability to differentiate into cells of all three embryonic germ layers and hence hold great promise for modeling and treating human diseases and conditions. However, during propagation in culture, hPSCs often become aneuploid with abnormal numbers of chromosomes (Baker et al., 2007; Mayshar et al., 2010; Taapken et al., 2011). Aneuploidy in hPSCs is attributed to culture adaptation that selects for abnormal, stable aneuploid karyotypes which outcompete diploid hPSCs limiting potential therapeutic applications (Baker et al., 2007; Keller and Spits, 2021; Mayshar et al., 2010; Price et al., 2021; Taapken et al., 2011).

Although culture adaptation explains how reoccurring constitutive aneuploidies become dominant in cultures of hPSCs, it does not explain how or why mitotic chromosome segregation errors occur in hPSCs generating an aneuploid genome. Perturbed DNA replication dynamics, DNA damage and defects in chromosome condensation that typically cause structural aneuploidies involving copy number alterations to chromosomal segments are linked to mitotic defects in hPSCs (Burrell et al., 2013; Halliwell et al., 2020; Lamm et al., 2016). However, whole chromosome aneuploidies resulting in the gain or loss of whole chromosomes are also prevalent in hPSCs (Baker et al., 2007; Mayshar et al., 2010; Taapken et al., 2011), but we do not know the mitotic pathways responsible for whole chromosome segregation errors in hPSCs.

Similarly, during early human embryogenesis aneuploidy is prevalent in totipotent and pluripotent embryonic cells with aneuploidy rates ranging between 25-90% for *in vitro fertilization* (IVF) preimplantation human embryos (Baart et al., 2006; Fragouli et al., 2008, 2013; McCoy et al., 2015; Mertzanidou et al., 2013; Vanneste et al., 2009) making aneuploidy the leading cause of miscarriages and birth defects in humans (Hassold and Hunt, 2001; Menasha et al., 2005; Orr et al., 2015). The high incidence of aneuploidy occurs irrespective of maternal age, infertility or embryo quality (Mertzanidou et al., 2013; Popovic et al., 2019; Vanneste et al., 2009). Due to obvious legal and ethical restrictions, aneuploidy rates in naturally conceived human embryos are unknown but are thought to correspond to aneuploidy rates in IVF preimplantation embryos accounting for a low human fecundity rate with only ∼30% of conceptions resulting in live births (Macklon et al., 2002; McCoy, 2017).

Surprisingly, like hPSCs, whole chromosome abnormalities caused by mitotic errors are more frequent than meiotic errors and structural aneuploidies in IVF preimplantation embryos (McCoy et al., 2015; Vanneste et al., 2009). This raises the intriguing possibility that mitotic errors and aneuploidy in hPSCs are not solely an artifact of growth in culture but rather are intrinsic characteristics of pluripotent cells. Yet, like hPSCs, we also do not know the underlying mechanisms causing the persistent and high rate of mitotic chromosome segregation errors in preimplantation embryonic cells. To investigate the mechanisms responsible for mitotic chromosome segregation errors in pluripotent cells, we categorize and quantify mitotic errors in hPSCs using fixed and time-lapse live-cell fluorescence microscopy. Furthermore, we use small molecules and manipulate development potential to test the influence of mitotic duration and chromosome microtubule attachment stability and developmental state, respectively, on chromosome segregation fidelity in hPSCs.

## Results

### Lagging chromosomes, caused by merotelic microtubule attachments, are responsible for elevated mitotic error rates in hPSCs

To determine the mechanisms causing mitotic chromosome segregation errors in hPSCs, we categorized the types of anaphase errors observed and compared anaphase error rates between pluripotent H1 and H9 human embryonic stem cells (hESCs) derived from the inner cell mass of human blastocysts (Thomson et al., 1998) and normal, primary somatic BJ fibroblasts (Figures 1A-B). Lagging chromosomes, unaligned chromosomes, and multipolar anaphases cause whole chromosome aneuploidy while acentric DNA fragments and chromosome bridges lead to structural aneuploidy (Figures 1A-B) (Burrell et al., 2013; Orr et al., 2015; Thompson and Compton, 2008). We also included a combination category for cells that exhibited multiple types of anaphase errors (Figures 1A-B).

**Figure 1.**
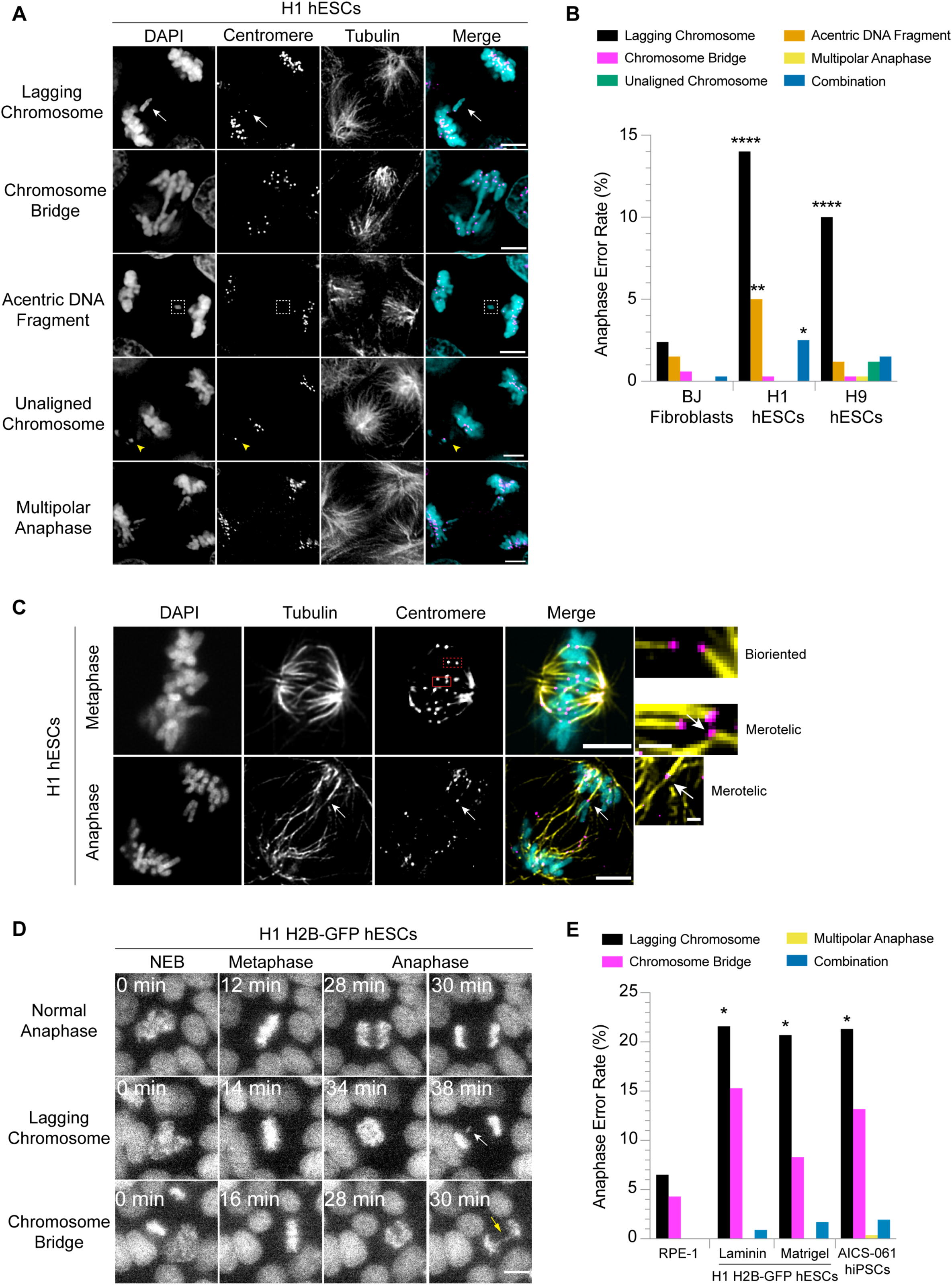
Mitotic error rates are elevated in hPSCs compared to somatic cells. (**A**) Representative images of anaphase errors including a lagging chromosome with a centromere (white arrow), chromosome bridge, acentric DNA fragment lacking a centromere (box), unaligned chromosome with a centromere (yellow arrowhead) and multipolar anaphase in H1 hESCs. Shown is DNA (cyan), centromeres (magenta) and microtubules (yellow). Scale bars: 5 mm. (**B**) Percentage of anaphase errors in primary somatic BJ fibroblasts and H1 and H9 hESCs plated as single cells on Laminin-521. n = 317 (BJ fibroblasts), 251 (H1 hESCs), and 283 (H9 hESCs) anaphases from three independent experiments; *p < 0.05, **p < 0.01, ****p < 0.0001 using a two-tailed Fisher’s exact test. (**C**) Representative images of chromosome microtubule attachments in metaphase and anaphase H1 hESCs. Shown is DNA (cyan), microtubules (yellow) and centromeres (magenta). In the metaphase cell, boxes are pairs of centromeres with a bioriented attachment (dashed) or a merotelic attachment (solid, white arrow). The anaphase cell shows a lagging chromosome with a merotelic attachment (white arrow). Insets are magnified views. Scale bars: 5 µm (main) and 1 µm (insets). (**D**) Selected panels from time-lapse live-cell fluorescence imaging of H1 H2B-GFP hESCs showing a normal anaphase and erroneous anaphases with a lagging chromosome (white arrow) or chromosome bridge (yellow arrow). Scale bar: 10 µm. (**E**) Percentage of anaphase errors in somatic RPE-1 H2B-GFP, H1 H2B-GFP hESCs plated as single cells on Laminin-521 or as aggregates on Matrigel and AICS-061 hiPSCs. n = 46 anaphases in RPE-1 and n = 111 (H1 on Laminin-521), 121 (H1 on Matrigel), and 258 (AICS-061) anaphases in hPSCs from at least three independent experiments; *p < 0.05 using a two-tailed Fisher’s exact test. See also Figure S1.

In somatic BJ fibroblasts, lagging chromosomes were the most frequent anaphase error; however, the rate was less than 5% (Figure 1B). In agreement, mitotic error and aneuploidy rates are less than 5% in other normal human somatic cells and tissues (Cimini et al., 1999; Knouse et al., 2014; Thompson and Compton, 2008). Lagging chromosomes were also the most frequent anaphase error in H1 and H9 hESCs; however, the rate was significantly higher, more than double (>10% in H1 and H9 hESCs), compared to somatic BJ fibroblasts (Figure 1B). Furthermore, H1 hESCs had a significantly higher frequency of acentric DNA fragments compared to somatic BJ fibroblasts. Though, it was less than half the frequency of lagging chromosomes in H1 hESCs, and there was not a similar trend in H9 hESCs (Figure 1B). In parallel samples, we quantified that more than 95% of the H1 or the H9 hESC population expressed the pluripotency transcription factors OCT4 or NANOG demonstrating that spontaneously differentiated cells did not account for the elevated error rates (Figures S1A-B).

As lagging chromosomes were the most frequent anaphase error in hESCs (Figure 1B), we sought to determine the causes of these. Using a calcium stable microtubule assay (Warren et al., 2020), we examined chromosome microtubule attachment orientations in hESCs because in other mammalian and human cancer cells, lagging chromosomes are caused by the persistence of improper merotelic chromosome microtubule attachments (also referred to as kinetochore microtubule or k-MT) in anaphase with a chromosome simultaneously attached to microtubules from both spindle poles (Cimini et al., 2001; Thompson and Compton, 2008; Thompson et al., 2010). In metaphase, H1 and H9 hESCs had correct bioriented attachments with sister chromatids attached to microtubules from opposite spindle poles and incorrect merotelic attachments with a single chromatid simultaneously attached to microtubules from both spindle poles (Figures 1C and S1C). Notably, 73% of the H1 and 50% of the H9 lagging chromosomes in anaphase had merotelic attachments (Figures 1C and S1C). We could not reliably conclude the attachment orientations of the remaining lagging chromosomes because in some instances, we observed a lagging chromosome attached to microtubules extending in opposite directions, but we could not track the microtubules back to the spindle poles. Also, the absence of a merotelic attachment may reflect the calcium sensitivity of the attachment rather than an alternative attachment orientation. Because of these limitations, our results likely underestimate the proportion of lagging chromosomes with merotelic attachments in hESCs.

HESCs often acquire stable chromosome abnormalities during culturing (Baker et al., 2007; Mayshar et al., 2010; Taapken et al., 2011) that, when coupled with the increased genomic instability caused by an aneuploid genome (Passerini et al., 2016; Sheltzer et al., 2011), suggests the possibility that only aneuploid hESCs exhibit erroneous lagging chromosomes with merotelic attachments. To address this possibility, prior to performing the calcium stable microtubule assay, we karyotyped both H1 and H9 hESC populations to monitor genomic stability since it is not feasible to simultaneously measure anaphase errors and determine the karyotype of cells. In both the H1 and the H9 populations, 20 of 20 cells scored were diploid. From this data, we can infer that less than 14% of cells in either population are aneuploid with 95% confidence (Baker et al., 2016) arguing that at least some H1 and H9 hESCs exhibiting lagging chromosomes with merotelic attachments are diploid.

To further validate our findings, we performed time-lapse live-cell fluorescence microscopy using normal, immortalized somatic RPE-1 H2B-GFP epithelial cells, H1 H2B-GFP hESCs (Calder et al., 2013), and AICS-061 human induced pluripotent stem cells (hiPSCs) that were derived from parental WTC-11 hiPSCs reprogrammed from dermal fibroblasts (Hayashi et al., 2016). Somatic RPE-1 H2B-GFP cells and H1 H2B-GFP hESCs exogenously express histone H2B tagged with GFP while AICS-061 hiPSCs express endogenous H2B monoallelically tagged with mEGFP allowing us to quantify anaphase error rates (Figures 1D-E and Videos S1-3). Similar to our previous results, lagging chromosomes were the most frequent anaphase error, and the rate was significantly elevated in H1 H2B-GFP hESCs (22% Laminin-521 and 21% Matrigel) and AICS-061 hiPSCs (21%) compared to somatic RPE-1 H2B-GFP cells (7%) (Figure 1E and Video S2). Although we cannot definitively distinguish acentric DNA fragments from lagging chromosomes in these experiments, we classified these errors as lagging chromosomes based upon the low incidence of acentrics in all our other analyses (Figures 1B and S3B, S4C and S4E). Thus, a shared phenotype of pluripotent cells is a high mitotic error rate compared to somatic cells with lagging chromosomes being the most frequent error. Furthermore, the lagging chromosome rate was significantly elevated in H1 H2B-GFP hESCs compared to somatic RPE-1 H2B-GFP cells irrespective of whether we dissociated and seeded H1 H2B-GFP hESCs as single cells without initial cell-cell contacts on a Laminin-521 substrate or as aggregates that maintain cell-cell contacts on a Matrigel substrate (Figure 1E). Our results combined with the high incidence of mitotic errors in IVF preimplantation embryos (McCoy et al., 2015; Vanneste et al., 2009), which maintain their 3D structure, argue that the disruption of tissue structure is unlikely to artificially increase mitotic error rates for hESCs growing in culture in contrast to recent findings in somatic epithelial tissues (Knouse et al., 2018).

Chromosome bridges were the second most frequent error, but there was not a significant difference in the chromosome bridge rate between somatic RPE-1 cells and hPSCs (Figure 1E). Also, we rarely observed multipolar anaphases or unaligned chromosomes (1/258 each in AICS-061 hiPSCs) (Figure 1E). The low incidence of unaligned chromosomes in hPSCs (Figures 1B and E) indicates the spindle assembly checkpoint (SAC) is functionally preventing anaphase onset until microtubules attach to chromosomes, which facilitates chromosome alignment (Musacchio and Salmon, 2007). In further support, we observed an H1 H2B-GFP hESC that delayed anaphase onset for more than 2 hrs due to a chromosome that failed to align (Figure S1D and Video S4), and hESCs arrest in mitosis in the presence of the microtubule depolymerizing drug nocodazole (Becker et al., 2006; Zhang et al., 2019). Thus, SAC signaling is functional and responsive to unattached chromosomes in hPSCs. In contrast, mouse morulae stage embryonic cells exhibit a high frequency of unaligned chromosomes indicative of insufficient SAC signaling (Vázquez-Diez et al., 2019). Collectively, our results agree with previous studies that quantified total mitotic error rates between 15-20% in hPSCs (Halliwell et al., 2020; Lamm et al., 2016; Milagre et al., 2020; Peterson et al., 2011; Taapken et al., 2011; Zhang et al., 2019), but importantly we extend these observations and demonstrate that lagging chromosomes in anaphase, caused by improper merotelic attachments, are the most frequent mitotic error in hPSCs and that lagging chromosome rates are significantly elevated in hPSCs compared to somatic cells.

### Prolonging mitotic duration decreases mitotic error rates in hPSCs

Chromosome missegregation rates are proportional to lagging chromosome rates because lagging chromosomes with merotelic attachments have an increased likelihood of segregating to the incorrect daughter cell producing two aneuploid progeny (Thompson and Compton, 2008). Therefore, we investigated why lagging chromosomes with merotelic attachments are more prevalent in hPSCs compared to somatic cells and on testing strategies to reduce merotelic attachments and decrease lagging chromosome rates in hPSCs, particularly because culture adaptation that selects for hPSCs with abnormal, stable aneuploid karyotypes must initiate with mitotic errors to generate the aneuploid substrates for selection (Baker et al., 2007; Taapken et al., 2011).

Improper merotelic attachments are not detected by the SAC (Cimini et al., 2001) but instead a network of kinases (Cimini et al., 2006; Godek et al., 2014; Salimian et al., 2011) and microtubule depolymerases (Bakhoum et al., 2008; Godek et al., 2014) converts improper merotelic attachments to correct bioriented attachments by facilitating iterative cycles of microtubule detachment and reattachment prior to anaphase onset (Godek et al., 2014). Thus, one parameter that influences merotelic error correction efficiency is mitotic duration with a longer mitotic duration allowing for more cycles of microtubule detachment and reattachment decreasing the frequency of errors (Figure 2A, note: error correction rate does not change) (Cimini et al., 2003; Sansregret et al., 2017) and conversely a shorter mitotic duration increasing errors. Accordingly, if mitotic duration is insufficient for robust merotelic error correction in hPSCs this will cause an elevated frequency of lagging chromosomes. A prediction of this hypothesis is that mitotic duration is shorter in hPSCs than somatic cells.

**Figure 2.**
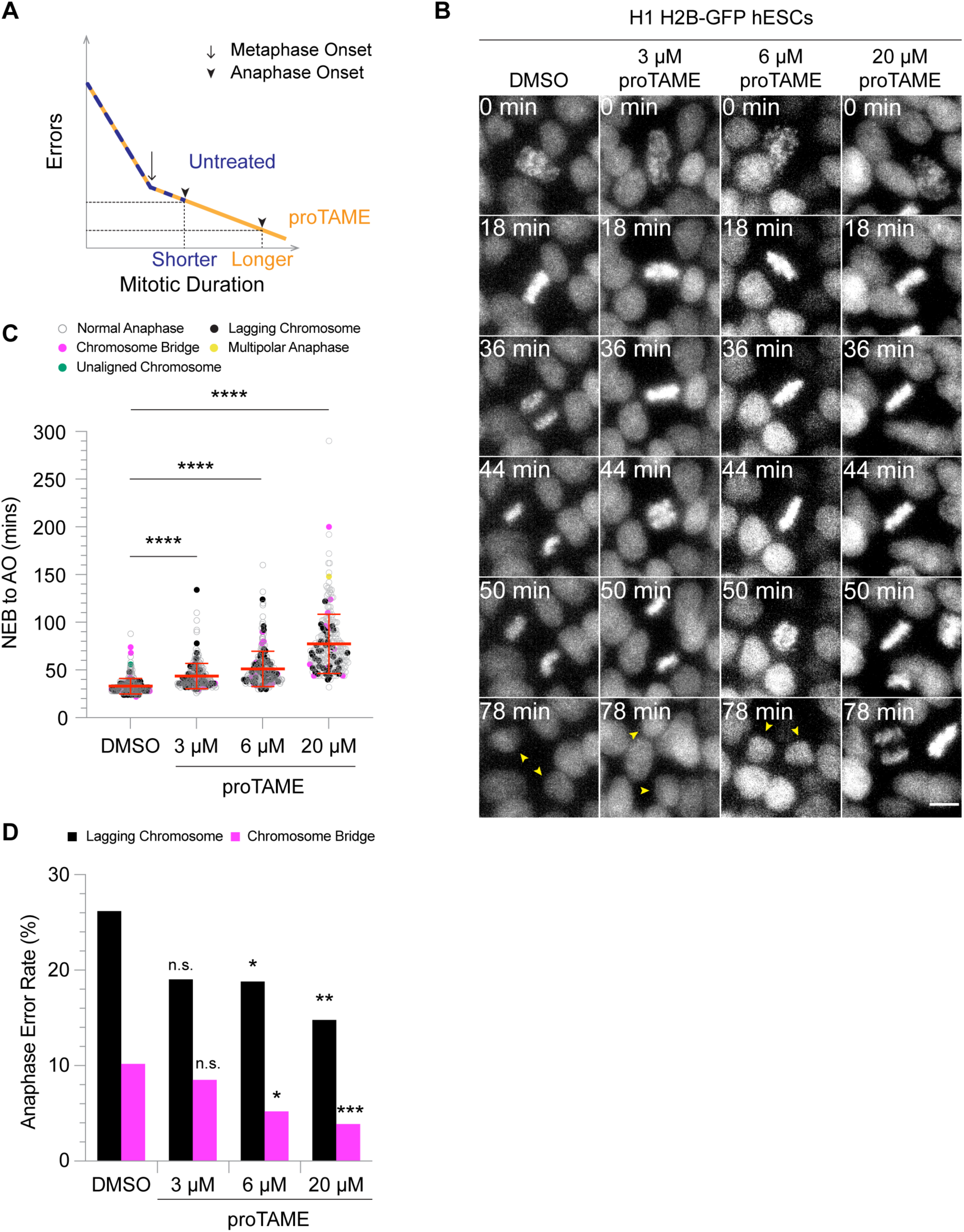
Prolonging mitotic duration decreases mitotic error rates in hPSCs. (**A**) Model illustrating the relationship between mitotic errors and mitotic duration. Early in mitosis improper chromosome microtubule attachments are prevalent due to the stochastic interaction of microtubules with chromosomes, but errors decline as mitosis progresses and improper attachments are converted to correct ones. Prolonging mitosis using the small molecule proTAME increases the amount of time for microtubule error correction reducing the frequency of mitotic errors. (**B**) Selected panels from time-lapse live-cell fluorescence imaging of H1 H2B-GFP hESCs that were treated with DMSO or increasing concentrations of proTAME (yellow arrowheads indicate daughter nuclei). Scale bar: 10 µm. (**C, D**) Mitotic duration (**C**) and percentage of lagging chromosomes or chromosome bridges (**D**) in H1 H2B-GFP hESCs treated with DMSO or increasing concentrations of proTAME. n = 275 (DMSO), 247 (3 µM proTAME), 250 (6 µM proTAME), and 257 (20 µM proTAME) anaphases from six independent experiments; NEB: nuclear envelope breakdown; AO: anaphase onset; mean ± SD and ****p < 0.0001 using a one-way ANOVA and Dunnett’s multiple comparisons test (**C**); n.s. p > 0.05, *p < 0.05, **p < 0.01, ***p < 0.001 using a two-tailed Fisher’s exact test (**C**). See also Figure S2.

To test this hypothesis, we measured mitotic duration from nuclear envelope breakdown (NEB) to anaphase onset (AO) in the H1 H2B-GFP hESCs, AICS-061 hiPSCs, and somatic RPE-1 H2B-GFP cells that we quantified anaphase error rates in (Figure 1E). In both H1 H2B-GFP hESCs and AICS-061 hiPSCs mitotic duration from NEB to AO, including prometaphase (NEB to metaphase) and metaphase (metaphase to AO), was significantly increased compared to somatic RPE-1 cells demonstrating that the elevated lagging chromosome rates in hPSCs are not caused by an abbreviated mitotic duration compared to somatic cells (Figure S1E). In further support, there was no significant difference in mitotic duration, including prometaphase or metaphase, between hPSCs that went through a normal mitosis or an aberrant mitosis (Figure S1F) underscoring that an abbreviated mitosis does not account for errors. These results suggest that other mechanisms are responsible for the elevated frequency of lagging chromosomes in hPSCs compared to somatic cells (see next section).

Nevertheless, we tested if prolonging mitosis would effectively reduce lagging chromosome rates in hPSCs by allowing for more cycles of microtubule release and reattachment prior to anaphase onset (Figure 2A). To test this strategy, we used the small molecule proTAME to delay mitotic progression. ProTAME inhibits the anaphase promoting complex/cyclosome (APC/C) E3 ubiquitin ligase whose activity is required for mitotic exit and whose partial inhibition increases mitotic duration in human somatic and cancer cells (Zeng et al., 2010). As a positive control, we reproduced previous results demonstrating that prolonging mitosis with proTAME reduces the frequency of mitotic errors, including lagging chromosomes, in somatic RPE-1 cells when error rates are artificially elevated (Figures S2A-C) (Sansregret et al., 2017). For our experiments in hPSCs, we added proTAME to H1 H2B-GFP hESCs or AICS-061 hiPSCs immediately prior to starting time-lapse live-cell imaging and imaged cells for 7 hrs in the presence of proTAME. Also, in parallel samples, we quantified that more than 95% of the H1 H2B-GFP or the AICS-061 hPSCs expressed OCT4 or NANOG prior to proTAME treatment, indicating that spontaneously differentiated cells in the populations were unlikely to influence the outcomes (Figures S2G and J).

In both H1 H2B-GFP hESCs and AICS-061 hiPSCs, mitotic duration significantly increased proportionally with proTAME concentration (Figures 2B-C and S2H). Importantly, as mitotic duration increased, the incidence of lagging chromosomes significantly decreased for H1 H2B-GFP hESCs (Figures 2C-D). The chromosome bridge and total anaphase error rates also significantly decreased with proTAME treatment (Figures 2C-D and S2E). Chromosome bridges are a consequence of under-replicated DNA regions or unresolved aberrant DNA structures that persist into mitosis and prolonging mitosis may also facilitate the correction of these errors (Fragkos and Naim, 2017). Surprisingly, the frequency of lagging chromosomes did not significantly decrease in AICS-061 hiPSCs (Figure S2I) despite an increased mitotic duration comparable to H1 H2B-GFP hESCs (Figures 2C and S2H). Consequently, we checked the genomic stability of AICS-061 hiPSCs during these experiments reasoning that aneuploid cells could be insensitive to this approach. There was a clonal abnormal karyotype, including a terminal deletion of the long arm of chromosome 18, but it was present at a low frequency in the population (10%, 2/20) and thus is unlikely to be the reason for the different response. We speculate that in AICS-061 hiPSCs other parameters have a greater influence on merotelic error correction. Also, we note that the lagging chromosome rate (∼20%) is approximately double the frequency of aneuploid cells in the population indicating that aneuploid cells do not solely account for the error rate.

The reduction in the lagging chromosome rate in H1 H2B-GFP hESCs could be a consequence of prolonging prometaphase, metaphase or both. Interestingly, metaphase was selectively lengthened proportional to proTAME concentration (Figure S2D) in H1 H2B-GFP hESCs, similar to somatic RPE-1 H2B-GFP cells (Figure S2A), demonstrating that, at least for some cells, metaphase duration can be a rate-limiting step in merotelic error correction. In further support, there was a significant decrease in metaphase duration for H1 H2B-GFP hESCs that went through mitosis with a lagging chromosome vs. a normal mitosis in the 20 µM proTAME treatment group (Figure S2F). However, this trend did not occur in the 6 µM proTAME treatment group (Figure S2F) revealing that error correction is not exclusively limited by metaphase duration. Combined, our results demonstrate that the elevated frequency of lagging chromosomes in hPSCs compared to somatic cells is not caused by an abbreviated mitosis; however, delaying mitotic progression, and metaphase specifically, is an effective strategy to improve merotelic error correction and reduce the lagging chromosome rate in hPSCs, albeit with the application limited to select hPSC lines.

### Decreasing microtubule attachment stability reduces mitotic errors in hPSCs

The iterative cycles of microtubule detachment and reattachment required for merotelic error correction also dictate that the error correction rate depends on chromosome microtubule attachment turnover with hyperstable attachments (i.e. low turnover) inhibiting the release of incorrect merotelic attachments (Bakhoum et al., 2009; Godek et al., 2014). Hence, hyperstable chromosome microtubule attachments in hPSCs relative to somatic cells is an alternative hypothesis explaining the elevated incidence of lagging chromosomes in hPSCs. This predicts that decreasing microtubule attachment stability (i.e. increasing turnover) will reduce lagging chromosome rates in hPSCs (Figure 3A, note: mitotic duration does not change).

**Figure 3.**
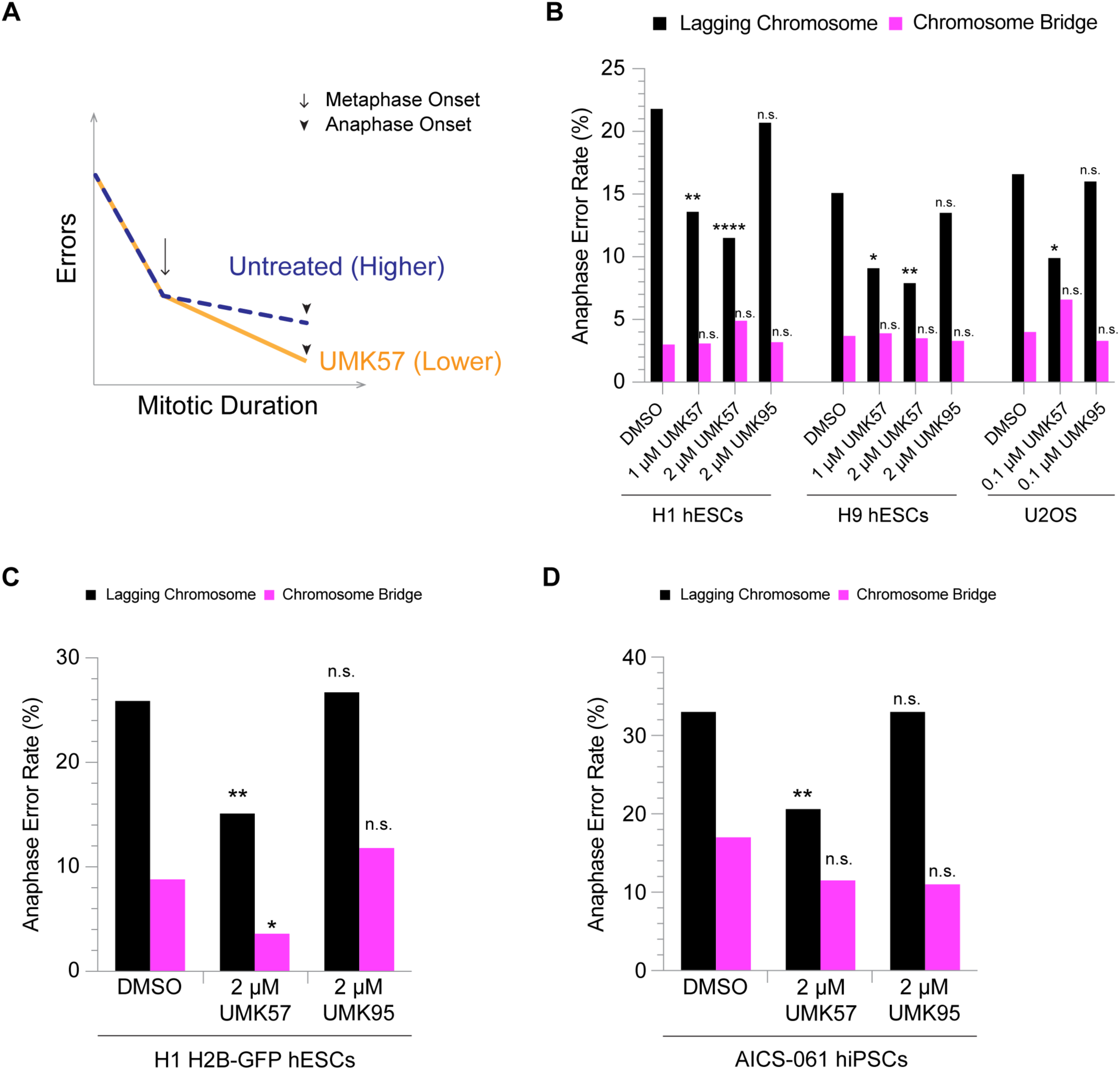
Decreasing microtubule attachment stability reduces mitotic errors in hPSCs. (**A**) Model illustrating the relationship between mitotic errors and chromosome microtubule attachment stability. Higher microtubule attachment stability decreases the correction rate of improper attachments while lowering microtubule stability using the small molecule UMK57 increases the correction rate of improper attachments reducing mitotic errors. (**B**) Percentage of lagging chromosomes and chromosome bridges in H1 and H9 hESCs and U2OS cancer cells after treatment with DMSO, UMK57 or UMK95 an inactive analog of UMK57 for 45 mins. n = 431 (DMSO), 390 (1 µM UMK57), 391 (2 µM UMK57), and 410 (2 µM UMK95) anaphases in H1 hESCs. n = 405 (DMSO), 385 (1 µM UMK57), 367 (2 µM UMK57), and 393 (2 µM UMK95) anaphases in H9 hESCs. n = 199 (DMSO), 242 (0.1 µM UMK57), and 243 (0.1 µM UMK95) anaphases in U2OS from three independent experiments; n.s. p > 0.05, *p < 0.05, **p < 0.01, ****p < 0.0001 using a two-tailed Fisher’s exact test. (**C, D**) Percentage of lagging chromosomes and chromosome bridges from time-lapse live-cell fluorescence imaging of H1 H2B-GFP hESCs (**C**) or AICS-061 hiPSCs (**D**) treated with DMSO, UMK57 or UMK95 for 12 hrs. n = 205 (H1, DMSO), 278 (H1, 2 µM UMK57), 187 (H1, 2 µM UMK95), 158 (AICS-061, DMSO), 209 (AICS-061, 2 µM UMK57), and 177 (AICS-061, 2 µM UMK95) anaphases from three independent experiments; n.s. p > 0.05, *p < 0.05, **p < 0.01 using a two-tailed Fisher’s exact test. See also Figure S3.

To test this prediction, we used the small molecule UMK57, an agonist of the microtubule depolymerase mitotic centromere-associated kinesin (MCAK or KIF2C), to decrease microtubule attachment stability in hPSCs. In human cancer cells with hyperstable attachments, short-term UMK57 treatment potentiates MCAK activity destabilizing microtubule attachments in metaphase and thus reduces the lagging chromosome rate (Orr et al., 2016). We treated H1 and H9 hESCs and positive control U2OS cancer cells for 45 mins with UMK57 prior to measuring anaphase error rates. To control for off-target effects, we also measured errors in cells treated for 45 mins with UMK95, an inactive analog of UMK57 (Orr et al., 2016).

As expected, short-term UMK57 treatment in U2OS cancer cells significantly reduced the lagging chromosome rate while UMK95 treatment did not (Figure 3B) (Orr et al., 2016). Likewise, lagging chromosome rates were significantly reduced by approximately 50% in H1 and H9 hESCs treated with UMK57, but at higher concentrations, while UMK95 treatment did not (Figure 3B). Also, lagging chromosome rates were selectively reduced while other anaphase error rates were not (Figure 3B and S3B), highlighting that the mechanisms responsible for different types of mitotic errors are distinct. Furthermore, unlike in cancer cells, high doses of UMK57 did not affect H1 or H9 hESC mitotic progression (Figure S3A) (Orr et al., 2016). As previous, we determined that spontaneously differentiated cells in the H1 and the H9 populations did not account for the error rates (Figure S3E). Moreover, we performed these experiments using the same batch of H1 and H9 hESCs that we karyotyped for the calcium stable microtubule assay and showed were diploid within the sensitivity range for the number of cells scored. Combined, these results demonstrate that destabilizing chromosome microtubule attachments in hPSCs increases the rate of merotelic error correction, reducing the frequency of lagging chromosomes (Figure 3A).

Although we modeled the effects of mitotic duration and chromosome microtubule attachment stability on merotelic error correction as two separate and independent pathways (Figures 2A and 3A), these may influence error correction in a dependent manner. To test this possibility, we simultaneously measured mitotic duration and errors in H1 H2B-GFP hESCs and AICS-061 hiPSCs by time-lapse live-cell fluorescence microscopy in the presence of UMK57 for 12 hrs (Figures 3C-D and S3C-D). In contrast to prolonging mitosis with proTAME (Figures S2I and J), destabilizing microtubule attachments with UMK57 significantly reduced lagging chromosome rates in both H1 H2B-GFP hESCs (Figure 3C) and AICS-061 hiPSCs (Figure 3D) while UMK95 did not. The chromosome bridge rate also significantly decreased in H1 H2B-GFP hESCs (Figure 3C), but this was not consistent in the AICS-061 hiPSCs (Figure 3D).

Interestingly, there was a significant increase in mitotic duration with UMK57 treatment, and specifically metaphase, for both H1 H2B-GFP hESCs and AICS-061 hiPSCs while UMK95 treatment did not significantly affect it (Figures S3C-D). However, for H1 H2B-GFP hESCs, the increase in metaphase duration was comparable to 3 µM proTAME treatment (metaphase mean = 15.8 mins DMSO vs. 27.3 mins 3 µM proTAME and metaphase mean = 12.2 mins DMSO vs. 20.4 mins UMK57), which did not significantly reduce the lagging chromosome rate (Figure 2D). For AICS-061 hiPSCs, no amount of delay in mitotic progression reduced the lagging chromosome rate (Figure S2I) suggesting that potentiating MCAK depolymerase activity predominantly enhances error correction by destabilizing microtubule attachments. Thus, mitotic duration and chromosome microtubule attachment stability are largely two independent parameters that influence merotelic error correction efficiency.

During these experiments, we also monitored the genomic stability of H1 H2B-GFP hESCs and AICS-061 hiPSCs. Similar to our previous analysis, there were clonal aneuploid cells with a terminal deletion of the long arm of chromosome 18 present in the AICS-061 population at a low frequency (<10%, 3/32). For the H1 H2B-GFP hESCs, initial karyotyping done after performing two complete experimental sets found 20 of 20 cells were diploid. Subsequent karyotyping, after the third experimental set, identified a fraction of abnormal cells with an interstitial duplication of the long arm of chromosome 20 in the population (25%, 5/20). Overall, the reduction in lagging chromosome rates upon UMK57 treatment is reproducible using multiple different hPSC lines arguing that the low incidence of aneuploid cells is unlikely to influence the outcomes. Collectively, these results support our hypothesis that hyperstable chromosome microtubule attachments contribute to the elevated frequency of erroneous lagging chromosomes in both hESCs and hiPSCs compared to somatic cells and that decreasing microtubule attachment stability is an effective strategy to reduce lagging chromosome rates in hPSCs.

### Developmental potential influences mitotic error rates

Chromosome segregation errors are rare in somatic cells (Figures 1B and D) (Cimini et al., 1999; Thompson and Compton, 2008), so it is widely assumed that a high fidelity of chromosome segregation is conserved in normal, non-transformed cells. Consequently, our repeated observations that mitotic error rates, and particularly lagging chromosome rates, are elevated in hPSCs compared to somatic cells (Figures 1B and D) coupled with the high mitotic error rates in preimplantation human embryos (McCoy et al., 2015; Vanneste et al., 2009) challenge this assumption. The opposing phenotypes of somatic vs. embryonic cells and hPSCs with respect to the frequency of mitotic errors led us to question whether a high error rate is an intrinsic and a cell autonomous trait linked to developmental state. This idea predicts that mitotic error rates and developmental potential correlate such that as developmental potential decreases mitotic error rates decrease and that as developmental potential increases so do mitotic error rates (Figure 4A). We tested this prediction using isogenic cells with different developmental states to eliminate genetic diversity as a confounding variable.

**Figure 4.**
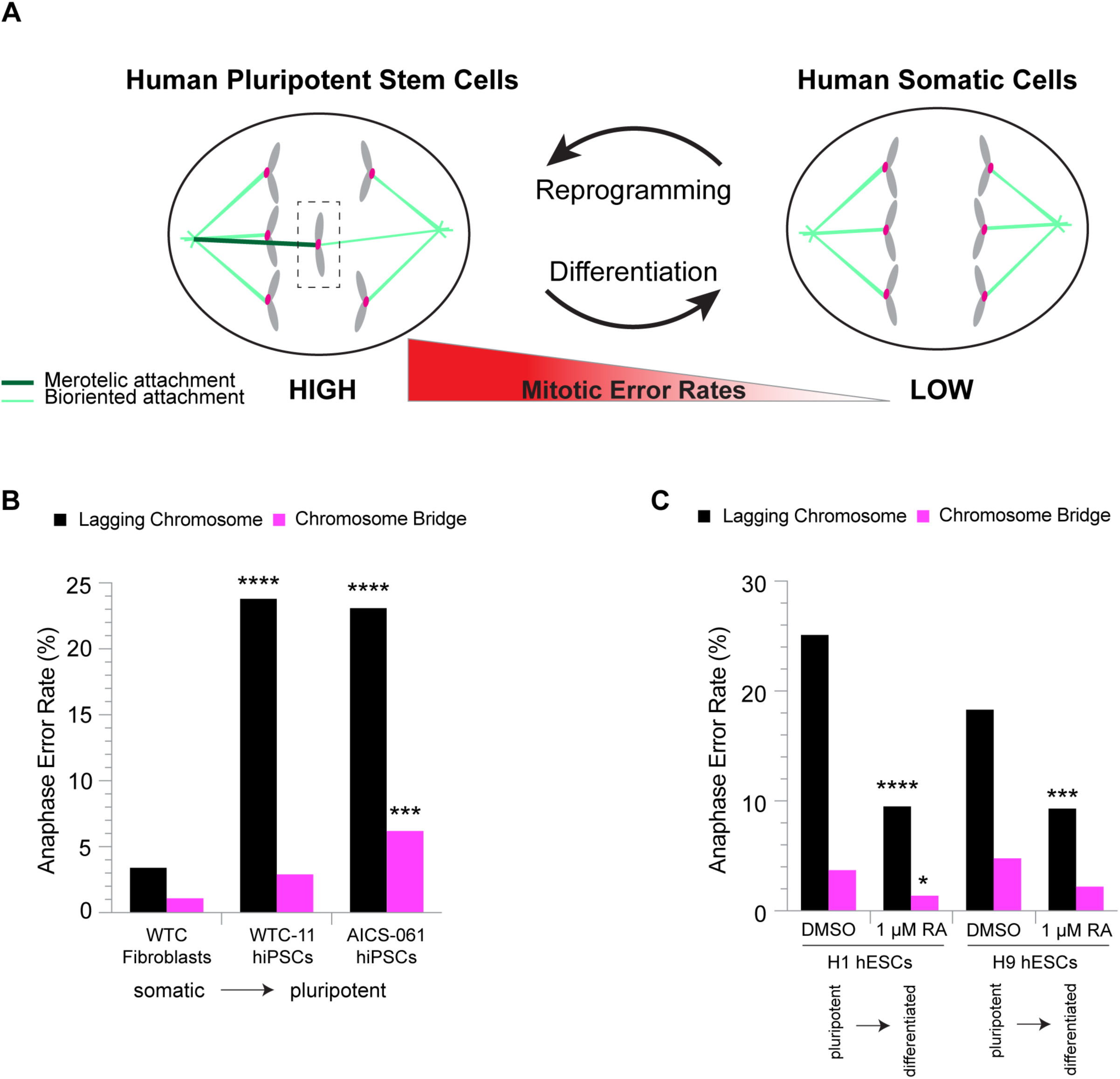
Developmental potential influences mitotic error rates. (**A**) Model illustrating the correlation between mitotic errors and developmental potential. As developmental potential decreases mitotic errors decrease, and conversely as developmental potential increases mitotic errors increase. (**B**) Percentage of lagging chromosomes and chromosome bridges in isogenic somatic WTC fibroblasts, WTC-11 hiPSCs and AICS-061 hiPSCs. n = 268 (WTC fibroblasts), 421 (WTC-11 hiPSCs), and 438 (AICS-061 hiPSCs) anaphases from three independent experiments; ***p < 0.001, ****p < 0.0001 using a two-tailed Fisher’s exact test. (**C**) Percentage of lagging chromosomes and chromosome bridges in H1 and H9 hESCs after 4 day treatment with DMSO or 1 µM all-*trans* retinoic acid (RA) to induce undirected differentiation. n = 454 (H1, DMSO), 358 (H1, 1 µM RA), 398 (H9, DMSO), and 356 (H9, 1 µM RA) anaphases from three independent experiments; *p < 0.05, ***p < 0.001, ****p < 0.0001 using a two-tailed Fisher’s exact test. See also Figures S4 and S5.

We compared mitotic error rates between isogenic normal, primary somatic WTC-11 fibroblasts to parental WTC-11 and the derivative AICS-061 hiPSCs. In agreement with our prediction, anaphase errors, with lagging chromosomes being the most frequent error, were significantly elevated in WTC-11 (lagging = 24%) and AICS-061 hiPSCs (lagging = 23%) compared to isogenic somatic WTC-11 fibroblasts (lagging = 3%) (Figure 4B and S4C). In addition, we karyotyped somatic WTC-11 fibroblasts and WTC-11 hiPSCs to confirm that abnormal aneuploid cells present in either population did not exclusively account for the error rates. Somatic WTC-11 fibroblasts were diploid (20/20) while 10% (2/20) of the WTC-11 hiPSCs had a clonal balanced translocation between the short arm of chromosome 1 and long arm of chromosome 16; however, even with the hypothetical assumption that all aneuploid cells go through an aberrant mitosis with a lagging chromosome and discarding 10% of the lagging chromosome data, lagging chromosome rates remained significantly elevated in WTC-11 hiPSCs compared to somatic WTC-11 fibroblasts (Figure S4D). Furthermore, to confirm the developmental states of isogenic WTC-11 and AICS-061 hiPSCs and somatic WTC-11 fibroblasts, we quantified the percent of cells expressing the pluripotency transcription factors OCT4 and NANOG. As expected, somatic WTC-11 fibroblasts did not express OCT4 and NANOG while nearly 100% of the WTC-11 and AICS-061 hiPSCs did (Figure S4B). Thus, with increased developmental potential mitotic error rates also increase.

If mitotic error rates correlate with developmental potential as we predict (Figure 4A), then differentiation and loss of pluripotency should decrease error rates. To test this, we induced undirected differentiation in H1 or H9 hESCs with *all-trans* retinoic acid (RA) (Jain et al., 2012). During a 4-day time course, DMSO treated control H1 and H9 hESCs maintained their pluripotent stem cell morphology of tightly packed colonies with smooth borders and a high nuclear to cytoplasmic ratio while RA treated hESCs acquired a flattened morphology and lower nuclear to cytoplasmic ratio (Figure S4F) indicative of differentiation. Also, expression of the pluripotency transcription factors OCT4, NANOG and SOX2 significantly decreased in the RA treated cells at the endpoint comparable to levels in somatic WTC-11 fibroblasts (Figures S5A-C) indicating loss of pluripotency. Importantly, after 4 days of RA undirected differentiation, anaphase error rates, including lagging chromosomes, were significantly decreased by approximately 50% compared to DMSO control H1 or H9 hESCs (Figures 4C and S4E) demonstrating that decreasing developmental potential reduces mitotic error rates. We also observed a slight, but significant increase, in multipolar anaphases; however, the frequency was less than 3% (Figure S4E).

In chimeric mouse embryos and human gastruloids composed of mixed populations of diploid and aneuploid cells, aneuploid cells are depleted as development progresses and differentiation occurs (Bolton et al., 2016; Yang et al., 2021). Analogous to this is the possibility that aneuploid cells present in the starting H1 and H9 populations used for the RA experiments are responsible for the mitotic errors but become depleted during differentiation thus decreasing the error rate. This scenario requires that H1 and H9 hESC populations are composed of aneuploid cells or are mosaic populations of diploid and aneuploid cells. Therefore, we karyotyped the H1 and the H9 hESC populations after completion of all experimental replicates reasoning that clonal and/or non-clonal aneuploidies were most likely to be detected after prolonged culturing. Critically, both the H1 and H9 populations were diploid (20/20) arguing that depletion of aneuploid cells during differentiation is unlikely to explain the decrease in anaphase errors. Collectively, our results show that mitotic error rates correlate with developmental potential and suggest that a high mitotic error rate is an inherent and cell autonomous trait of hPSCs.

## Discussion

Here we show that lagging chromosomes in anaphase, caused by persistent improper merotelic chromosome microtubule attachments, are the most frequent mitotic error in hPSCs. Surprisingly, our results reveal that hPSCs are more similar to transformed human cancer cells than non-transformed normal somatic cells with respect to mitotic error rates, particularly lagging chromosome rates (Cimini et al., 2001; Godek et al., 2016; Thompson and Compton, 2008). Furthermore, we show that mitotic error rates correlate with developmental potential decreasing upon loss and increasing upon gain, demonstrating that a high mitotic error rate is intrinsic to hPSCs. In agreement, multipotent neural stem cells exhibit an intermediate error rate (∼10%) between hPSCs and somatic cells suggesting a linear correlation with developmental potential (Godek et al., 2016). Collectively, these results demonstrate that a high fidelity of chromosome segregation is not universally conserved in normal, diploid human cells and that it depends on developmental state. This raises the possibility that in cancer cells the (re)acquisition of a developmental program with greater potency rather than of mutations in mitotic genes causes an elevated mitotic error rate in agreement with the low frequency of genetic alterations found in mitotic genes (Greenman et al., 2007; Nath et al., 2015).

Assuming that the chromosome missegregation and lagging chromosome rates are proportional in hPSCs, analogous to cancer cells (Thompson and Compton, 2008), then lagging chromosomes are a leading cause of aneuploidy in hPSCs. In hPSCs, the ∼20% lagging chromosome rate is comparable to that of HT29 colon cancer cells which corresponds to a ∼0.3% missegregation rate per chromosome (Thompson and Compton, 2008). Using this benchmark, we estimate that hPSCs missegregate a chromosome every tenth division. We assume that missegregation would be random as there is no known bias to preferentially missegregate a chromosome in unperturbed conditions. Ideally, we would directly measure chromosome missegregation rates, but the growth of hPSCs as tightly packed colonies combined with their poor survival as single cells poses challenges to using conventional techniques (Godek and Compton, 2018; Thompson and Compton, 2008).

Although we estimate a high chromosome missegregation rate, we detect a low frequency of aneuploid hPSCs in culture, indicating that most aneuploid hPSCs are at a selective disadvantage, thus maintaining a predominately diploid population. In this regard, hPSCs resemble somatic cells which arrest in the subsequent cell cycle following chromosome missegregation preserving a homogeneous diploid karyotype (Thompson and Compton, 2010). In contrast, cancer cells tolerate and propagate with aneuploid genomes (Godek et al., 2016; Thompson and Compton, 2010). Alternatively, our estimate may be an overestimate, and the generation of aneuploid progeny is a rarer event in hPSCs. Regardless of the exact missegregation rate, these results delineate a pathway driving the process of culture adaptation in hPSCs that selects for reoccurring stable chromosome abnormalities which do outcompete diploid hPSCs (Baker et al., 2007; Mayshar et al., 2010; Taapken et al., 2011). We propose the process depends on a lagging chromosome that leads to chromosome missegregation producing aneuploid progeny which are then substrates for culture selection pressures to act on (Figure 5A). In this scenario, lagging chromosomes, which are an inherent and cell autonomous trait, are the key agents of change fueling culture adaptation, but this must also be coupled to the transient survival of aneuploid hPSCs providing an opportunity for selection to occur. How hPSCs gain initial or transient tolerance to an aneuploid genome is unknown, but hPSCs often acquire p53 mutations (Merkle et al., 2017) and this may lead to aneuploidy tolerance as shown in cancer cells (Thompson and Compton, 2010). Subsequently, selection for aneuploid hPSCs with constitutive stable chromosome abnormalities that support long-term survival and propagation with a growth advantage over diploid hPSCs occurs (Price et al., 2021). This multi-step process also explains why culture adaptation often arises during extended culturing (Baker et al., 2007).

**Figure 5.**
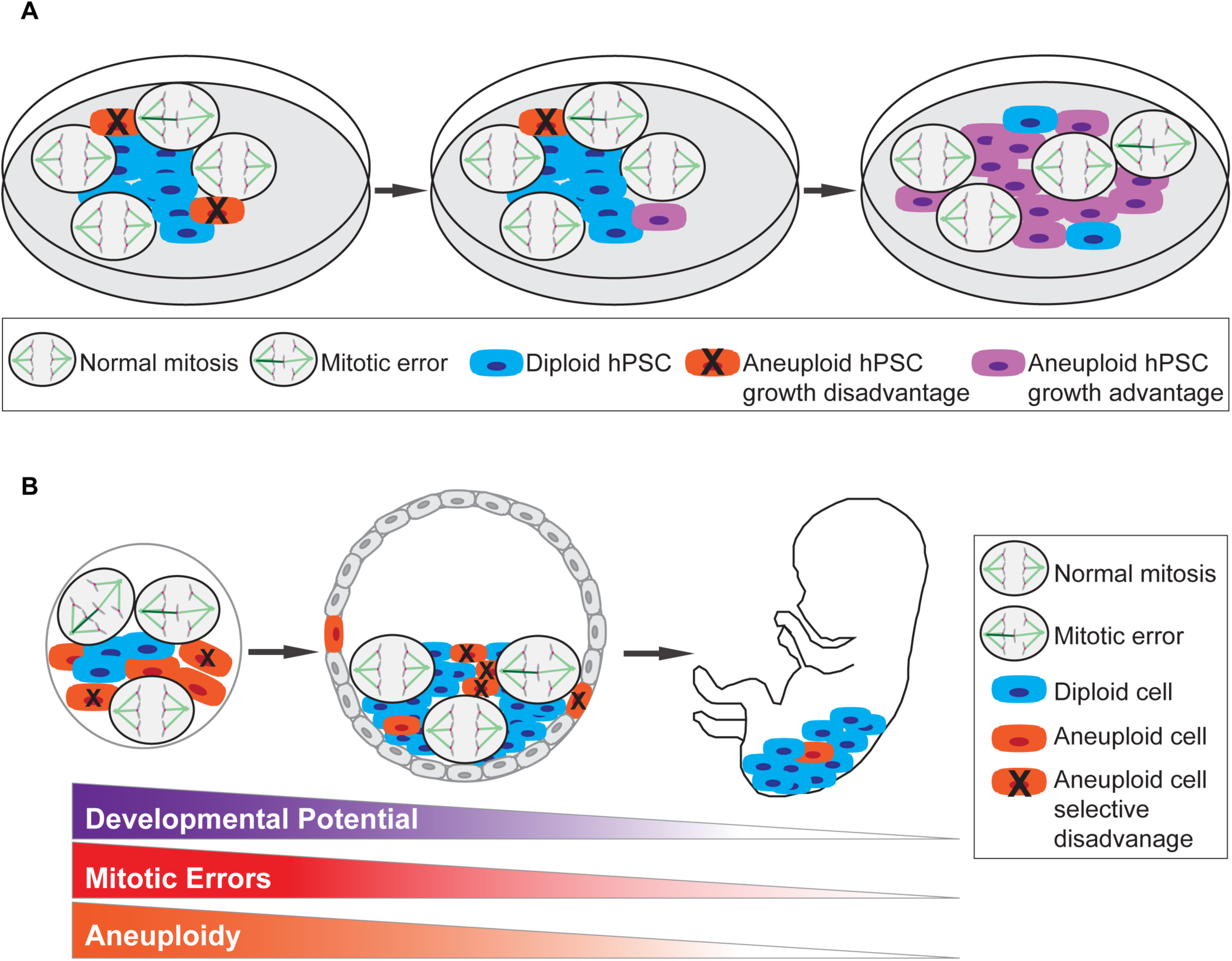
Models for how mitotic errors contribute to culture adaptation in hPSCs and aneuploidy during human development. (**A**) We speculate that culture adaptation in hPSCs depends on lagging chromosome errors that lead to chromosome missegregations and the generation of aneuploid progeny. Most aneuploid hPSCs (orange) are at a growth disadvantage and are outcompeted by diploid hPSCs (blue) as chromosome missegregation is random with respect to which chromosome is missegregated. However, the persistent and high rate of lagging chromosome errors in hPSCs coupled with continued propagation in culture increases the probability that aneuploid hPSCs with stable chromosome abnormalities conferring a growth advantage over diploid hPSCs are selected for (purple hPSCs). (**B**) We propose that preimplantation totipotent and pluripotent embryonic cells exhibit a high rate of lagging chromosomes that cause chromosome segregation errors and the generation of aneuploid embryonic cells leading to mosaic embryos composed of mixed populations of diploid and aneuploid embryonic cells. However, as development progresses, developmental potential decreases coinciding with a decline in the lagging chromosome rate that when coupled with a selective disadvantage for aneuploid (orange) compared to diploid (blue) embryonic cells explains how mosaic embryos can support normal human development.

Given the causal relationship between lagging chromosomes and chromosome missegregation combined with the potential consequences of generating aneuploid progeny, understanding why merotelic attachments persist in hPSCs and devising strategies to reduce merotelic errors is paramount for the successful use of hPSCs in regenerative medicine therapies. Here we find that prolonging mitosis or destabilizing chromosome microtubule attachments using the small molecules proTAME or UMK57, respectively, improves merotelic error correction reducing lagging chromosomes in hPSCs. We note that prolonging mitosis using proTAME also decreases the incidence of unaligned chromosomes during mouse preimplantation development presumably by increasing attachment formation rather than merotelic error correction (Vázquez-Diez et al., 2019), suggesting that this strategy is broadly applicable. By extension we predict that these strategies should also suppress aneuploidy rates in hPSCs, although this remains to be tested. Of interest will be to test long-term UMK57 treatment in hPSCs as cancer cells, but not normal dermal fibroblasts (Barroso Vilares et al., 2020), become resistant to treatment (Orr et al., 2016). Also, it remains unknown if hPSCs maintain pluripotency during long-term treatment with these small molecules.

Furthermore, our UMK57 results suggest that, similar to cancer cells (Bakhoum et al., 2009), hyperstable microtubule attachments underlie the elevated frequency of lagging chromosomes in hPSCs. Measurement of microtubule attachment turnover rates in hPSCs will be necessary to test this. Although many molecular players regulating microtubule dynamics are known (Godek et al., 2014), how these networks differ between somatic cells and cancer cells or hPSCs is unknown. In contrast to aneuploid cancer cells where genetic and transcriptional heterogeneity is a confounding variable (Stingele et al., 2012; Zhao et al., 2019), hPSCs may offer a more tractable system to determine the molecular pathways causing hyperstable microtubule attachments as lagging chromosomes are not exclusive to aneuploid hPSCs.

Extending our results to human preimplantation development suggests that lagging chromosomes are primarily responsible for the high mitotic error and aneuploidy rates of early human embryonic cells (Figure 5B). In contrast, during mouse preimplantation development, unaligned chromosomes are the most frequent mitotic error (Vázquez-Diez et al., 2019) suggesting different mechanisms are responsible for chromosome missegregation in mouse vs. human embryogenesis. This difference may contribute to the discrepancy in aneuploidy rates with 5% of mouse embryos (Hassold and Hunt, 2001; Lightfoot et al., 2006; Wei et al., 2011) and 25-90% of human embryos exhibiting aneuploidy (Baart et al., 2006; Fragouli et al., 2008, 2013; McCoy et al., 2015; Mertzanidou et al., 2013; Vanneste et al., 2009). In addition, IVF preimplantation embryos exhibit the related phenomena of chromosomal instability (CIN) that requires (1) persistent chromosome missegregation coupled with (2) the survival and propagation of aneuploid progeny (Orr et al., 2015; Thompson and Compton, 2008, 2010) producing heterogeneous aneuploid cells in a single embryo (Mertzanidou et al., 2013; Vanneste et al., 2009). Although a recent study of preimplantation bovine embryos, models for human embryogenesis, found that a failure of parental pronuclei to properly cluster and condense their chromosomes led to an increase in errors (Cavazza et al., 2021), parental genome clustering is unique to the first mitotic division and thus cannot account for the repeated mitotic errors that must occur to generate embryos with a CIN phenotype. Rather, erroneous lagging chromosomes are not restricted to specialized mitotic divisions and thus provide a mechanism for the CIN phenotype of human preimplantation embryos (Mertzanidou et al., 2013; Vanneste et al., 2009). Furthermore, although we estimate that every tenth division in hPSCs generates aneuploid progeny, this may underestimate the chromosome missegegration rate in human embryos given the prevalence of CIN in IVF cleavage stage embryos indicating repeated mitotic errors occurring within a few divisions (Mertzanidou et al., 2013; Vanneste et al., 2009). Future investigations, using other model systems for human preimplantation development, will be necessary to determine if the same mechanisms are responsible for mitotic errors as in hPSCs.

The CIN phenotype of preimplantation embryos also requires at least an initial tolerance to an aneuploid genome. How this occurs and whether a similar mechanism supports a limited tolerance to an aneuploid genome in hPSCs (providing an opportunity for culture selection to occur) is unknown. Accordingly, this raises the question of how euploid embryos are established to support normal development. Like most aneuploid hPSCs, aneuploid preimplantation embryonic cells may be at a selective disadvantage when in competition with diploid embryonic cells. In support, some mosaic blastocysts composed of diploid and aneuploid cells were euploid 12 days post-fertilization (Popovic et al., 2019) and transferred mosaic IVF embryos can result in normal development and live births (Yang et al., 2021). Importantly, our results suggest that the establishment of euploid embryos is also supported by declining mitotic error rates as developmental potential decreases and differentiation occurs (Figure 5B). Thus, during human development genome stability is achieved because the time window comprising embryonic cells with high developmental potency and high mitotic error rates is limited. In contrast, the time window is unlimited for hPSCs growing in culture. In conclusion, we propose that in normal human cells developmental state differentially influences the fidelity of chromosome segregation and the response to aneuploidy.

## Experimental Procedures

### Cell Lines

Primary BJ fibroblasts (CRL-2522) and U2OS (HTB-96) cell lines used in this study are available from the American Type Culture Collection (ATCC). We generated RPE-1 cells stably expressing H2B-GFP using parental RPE-1 (CRL-4000) cells available from ATCC. H1/WA01 and H9/WA09 hESCs are available from WiCell Research Institute. WTC-11 (GM25256) and AICS-061 hiPSCs are available from the Coriell Institute for Medical Research and Allen Institute for Cell Science, respectively. H1 H2B-GFP hESCs used in this study were obtained from Dr. Jonathan S. Draper, McMaster University. H1 H2B-GFP hESCs also express a G1 reporter, but we did not monitor G1 phase in our experiments. WTC fibroblasts were obtained the Gladstone Stem Cell Core.

### Cell Culture

U2OS (XX) cells were grown in Dulbecco’s Modified Eagle’s Medium (DMEM) supplemented with 10% fetal calf serum (FCS), 50 U/mL penicillin and 50 µg/mL streptomycin and 250 µg/L Amphotericin B. RPE-1 H2B-GFP (XX) cells were grown in DMEM supplemented with 10% fetal calf serum (FCS), 50 U/mL penicillin and 50 µg/mL streptomycin, 250 µg/L Amphotericin B, 20 mM HEPES and 5 µg/ml blasticidin. BJ fibroblast (XY) cells were grown in Eagle’s Minimum Essential Medium (EMEM) supplemented with 10% fetal bovine serum (FBS) and 100 U/mL penicillin and 100 µg/mL streptomycin. WTC fibroblast (XY) cells were grown in DMEM supplemented with 10% FBS, 2 mM GlutaMAX-1 (ThermoFisher #35050061), 0.1 mM MEM nonessential amino acids and 100 U/mL penicillin and 100 µg/mL streptomycin. H1/WA01 hESCs (XY), H9/WA09 hESCs (XX), WTC-11 hiPSCs (XY) and AICS-061 hiPSCs (XY) were grown in mTeSR1 medium (StemCell Technologies #85870). H1 H2B-GFP hESCs (XY) were grown in mTeSR1 supplemented with 1 µg/mL puromycin. All pluripotent stem cell lines were routinely grown on hESC qualified Matrigel (Corning #354277). For routine passaging, H1, H9 and H1 H2B-GFP hESCs and WTC-11 hiPSCs were dissociated using versene according to WiCell or Coriell Institute protocols, respectively. AICS-061 hiPSCs were passaged using StemPro Accutase (ThermoFisher #A1110501) in the presence of ROCK inhibitor Y-27632 (Tocris #1254) for an initial ∼20 hrs according to Allen Institute protocols. WTC-11 fibroblasts were routinely passaged using TrypLE Select (ThermoFisher #12563011), and BJ fibroblasts, U2OS and RPE-1 H2B-GFP cells were passaged using 0.05% trypsin. All cell lines were validated as mycoplasma free (Sigma-Aldrich Lookout® Mycoplasma PCR Detection Kit # MP0035) and grown at 37°C in a humidified atmosphere with 5% CO_2_. The karyotypes of human pluripotent stem cell lines and isogenic WTC fibroblasts used in this manuscript were verified with by G-banded karyotyping provided by WiCell Research Institute.

RPE-1 H2B-GFP cells were transfected with the pBOS H2B-GFP vector (BD Biosciences) using Fugene 6 (Promega #E2691) following manufacturer’s instructions. RPE-1 cells stably expressing H2B-GFP were selected using 5 µg/ml blasticidin and subsequently single cell clones were isolated using limiting dilution.

### Immunofluorescence

H1 and H9 hESCs and WTC-11 hiPSCs were plated as aggregates on Matrigel-coated 18 mm glass coverslips in 12-well cell culture plates unless otherwise noted in the figure legends. Alternatively, H1 and H9 hESCs were dissociated to single cells using TrypLE Select and plated on Laminin-521 (Biological Industries #05-753-1F) coverslips coated at 0.5 µg/cm^2^. AICS-061 hiPSCs were dissociated to single cells using StemPro Accutase and plated on Matrigel-coated 18 mm glass coverslips in 12-well cell culture plates with ROCK inhibitor for an initial ∼20 hrs and then subsequently the ROCK inhibitor was washed out. For AICS-061 hPSCs, all experiments were performed at least 24 hrs after the removal of ROCK inhibitor. BJ fibroblasts, WTC fibroblasts and U2OS cells were plated on standard 18 mm glass coverslips in 12-well cell culture plates prior to fixation.

For quantification of the pluripotency transcription factors OCT4 and NANOG, cells were fixed with 3.5% paraformaldehyde for 5 mins at room temperature, permeabilized with Tris-buffered saline (TBS) with 0.1% Triton X-100 for 2 × 5 mins and blocked with TBS with 2% bovine serum albumin (BSA) and 0.1% Triton X-100 for 30 mins at room temperature or overnight at 4°C. Primary antibodies were diluted in TBS + 2% BSA + 0.1% Triton X-100 at 5 µg/mL (rabbit anti-OCT4, Abcam #ab19857), 1:200 (mouse anti-NANOG, Abcam #ab173368) or 1:150 (rabbit anti-NANOG, Abcam #ab109250), and coverslips were incubated for 2 hrs at room temperature. Cells were then washed with TBS + 2% BSA + 0.1% Triton X-100 for 4 × 5 mins. Secondary antibodies were diluted in TBS + 2% BSA + 0.1% Triton X-100 + 0.5 µg/mL DAPI at 1:1000 and coverslips were incubated for 1 h at room temperature. Cells were washed with TBS + 2% BSA + 0.1% Triton X-100 for 2 × 5 mins, TBS + 0.1% Triton X-100 for 5 mins and TBS buffer for 5 mins sequentially. Coverslips were mounted on glass slides using ProLong Gold antifade (ThermoFisher #P36934) reagent. For SOX2 immunofluorescence, cells were fixed in 3.5% paraformaldehyde for 5 mins, washed 2 × 5 mins with TBS + 0.1% Triton X-100, blocked with TBS + 2% BSA + 0.1% Triton X-100 + 10% donkey serum overnight at 4°C. Subsequently the standard immunostaining protocol as described above was followed. The SOX2 primary antibody was used at 10 µg/mL (mouse anti-SOX2, R&D Systems #MAB2018).

For measuring the frequency of chromosome segregation errors, cells were fixed with ice-cold methanol for 5 mins and then permeabilized with high-salt TBS (containing 225 mM NaCl) with 0.1% Triton X-100 for 2 × 5 mins and blocked with high-salt TBS with 2% BSA and 0.1% Triton X-100 for 30 mins at room temperature or overnight at 4°C. Primary antibodies were diluted in high-salt TBS + 2% BSA + 0.1% Triton X-100 at 1:4000 (mouse anti-a-tubulin, Sigma #T6199) and 2 µg/mL (rabbit anti-CENP-A, Dr. A. Straight. Stanford University). To assess calcium stable chromosome microtubule attachments, cells were pre-extracted with calcium buffer (100 mM PIPES, 1 mM MgCl_2_, 0.1 mM CaCl_2_, 1% Triton X-100, pH = 6.8) for 5 mins and subsequently fixed with 1% glutaraldehyde in PBS for 10 mins. Coverslips were washed with 0.5 mg/mL sodium borohydride (NaBH4, dissolved in PBS) for 2 × 10 mins and then rinsed with PBS prior to blocking with TBS + 2% BSA + 0.5% Triton X-100 for 30 mins at room temperature. Cells were stained with primary antibodies diluted with TBS + 2% BSA + 0.1% Triton X-100 at 1:1000 (human anti-ACA, Geisel School of Medicine at Dartmouth) and 1:4000 (mouse anti-α-tubulin, Sigma) following the standard immunostaining protocol as described above. The following secondary antibodies (diluted at 1:1000) were used in this study: donkey anti-mouse Alexa Fluor 488, goat anti-rabbit Alexa Fluor 594, donkey anti-mouse Alexa Fluor 647, donkey goat anti-human Alexa Fluor 594, donkey anti-rabbit Alexa Fluor 647 (ThermoFisher #A-21202, #A-11037, #A-31571, #A-11014 and #A-31573, respectively).

### Microscopy for Immunofluorescence

Images were acquired with either a Hamamatsu ORCA-Fusion Gen III Scientific CMOS camera mounted on a Nikon Eclipse Ti2E microscope with a Nikon CFI Plan Apo Lambda 60×, 1.4 numerical aperture oil immersion objective, an Andor cooled CCD camera mounted on a Nikon Ti microscope with a Nikon Plan Apo VC 60×, 1.4 numerical aperture oil immersion objective or a spinning-disc confocal microscopy system (Micro Video Instruments) featuring a Nikon Eclipse Ti microscope equipped with an Andor CSU-W1 two-camera spinning disc module, Andor dual Zyla sCMOS cameras, an Andor ILE laser module, and a Nikon Plan Apo Lambda 60×, 1.4 numerical aperture oil immersion objective at room temperature. Image series in the *Z*-axis were obtained using either 0.2 µm or 0.5 µm optical sections.

For experiments comparing the percentage of cells expressing a protein of interest or quantifications of proteins levels, images for each cell line were acquired with the same acquisition parameters and exposure times. Image deconvolution and contrast enhancement were performed using NIS Batch Deconvolution (Nikon), NIS Elements (Nikon), ImageJ (NIH) and Photoshop (Adobe). Images shown are maximum intensity projections (chromosome segregation errors) or sum intensity projections (chromosome microtubule attachments and pluripotency markers) of selected *Z*-planes.

Criteria for scoring chromosome segregation errors is as follows: the presence of a chromosome that lags behind the segregating chromosomal mass and has clear centromere staining in anaphase is scored as a lagging chromosome. The presence of a chromosome without centromere staining between two segregating chromosomal masses in anaphase is scored as an acentric DNA fragment. The presence of chromosome spanning between segregating chromosomal masses in anaphase is scored as a chromosome bridge. A chromosome that never aligns to the metaphase plane and presents proximal to the spindle pole at anaphase onset is scored as an unaligned chromosome. An anaphase where chromosomes segregate to more than two poles is scored as a multipolar anaphase. An anaphase with multiple errors were scored as combination.

### Quantification of Protein Expression

To determine the percentage of cells expressing a protein of interest, sum intensity projections were compiled from *Z*-stack images using ImageJ (NIH). Then the maximum and minimum display values were scaled equivalently among different cell lines using somatic cell lines as a negative background control. Single cells were then categorized as positive or negative for expression of a protein of interest. For quantification of protein levels, sum intensity projections were compiled from *Z*-stack images using ImageJ (NIH). Nuclei were randomly picked per field of view based upon the DAPI signal. An elliptical region of interest (ROI) was drawn to encompass the whole nucleus and then a slightly larger elliptical ROI was drawn to encompass both the nucleus and the background. The mean background intensity was calculated based on the in-between background region of the two ROIs. Expression level of a protein of interest in each nucleus was represented by the background subtracted mean intensity of the ROI that encompasses the nucleus.

### Time-Lapse Live-Cell Fluorescence Imaging

RPE-1 H2B-GFP cells were plated in standard DMEM media on 18 mm glass coverslips in 12-well cell culture plates and incubated overnight at 37°C in a humidified atmosphere with 5% CO_2_. The next day, coverslips were washed with phenol-free media supplemented with 0.1% DMSO, 3 µM or 6 µM proTAME (Tocris #I-440-01M) and mounted in modified rose chambers. For monastrol arrest and release experiments, the next day following overnight incubation, coverslips were washed into standard DMEM media supplemented with 100 µM monastrol (Tocris #1305) and maintained for 6 hrs at 37°C in a humidified atmosphere with 5% CO_2_. After 6 hrs, cells were released by washing into phenol-free media supplemented with 0.1% DMSO, 3 µM or 6 µM proTAME (Tocris) and mounted in modified rose chambers. Live-cell imaging was performed at 37°C using an Andor cooled CCD camera mounted on a Nikon Ti microscope with a Nikon Plan Apo VC 60×, 1.4 numerical aperture oil immersion objective with binning set to 2×2. Image series in the *Z*-axis were obtained using 1 µm optical sections. Cells were imaged for 16 hrs with a 2 min time interval for proTAME only experiments or 5 hrs with a 2 min time interval for monastrol arrest and release experiments

H1 H2B-GFP hESCs and AICS-061 hiPSCs were plated in standard mTeSR1 media on the 35 mm glass bottom dishes (MatTek #P35G-1.5-14-C) coated with Matrigel or Laminin-521 and incubated for 1-3 days at 37°C in a humidified atmosphere with 5% CO_2_ prior to live-cell imaging. HPSCs were washed with phenol-free mTeSR1 media three times to get rid of spent media and then cultured in phenol-free mTeSR1 during live-cell imaging. Live-cell imaging was performed at 37°C in a humidified environment with 5% CO_2_ (Tokai Hit Stage-top Incubation System) using either a Hamamatsu ORCA-Fusion Gen III Scientific CMOS camera mounted on a Nikon Eclipse Ti2E microscope with a Nikon CFI Plan Apo Lambda 60×, 1.4 numerical aperture oil immersion objective with binning set to 2×2 or using spinning-disc confocal microscopy system (Micro Video Instruments) featuring a Nikon Eclipse Ti microscope equipped with an Andor CSU-W1 two-camera spinning disc module, Andor dual Zyla sCMOS cameras, an Andor ILE laser module, and a Nikon Plan Apo Lambda 60×, 1.4 numerical aperture oil immersion objective with binning set to 2×2. Image series in the *Z*-axis were obtained using 1 µm optical sections. HPSCs were imaged for 7 hours with 2 min time interval (wide-field fluorescence) or 12 hours with 2 min time interval (spinning-disc confocal). Of note, we optimized these experiments using the lowest exposure and intensity settings permissible to visualize errors while minimizing artifacts due to phototoxicity.

Image acquisitions and analyses were performed using NIS Elements (Nikon) and ImageJ (NIH). Representative images from live-cell imaging shown in this study are maximum intensity projections of all *Z-*planes or a single *Z*-plane. Cells undergoing mitosis were tracked from nuclear envelope breakdown (NEB) to anaphase onset, during which prometaphase (NEB to metaphase), metaphase (metaphase to anaphase onset) or total mitotic (NEB to anaphase onset) durations were recorded. In combination with mitotic duration, anaphase errors including lagging chromosomes, chromosome bridges, multipolar anaphases, unaligned chromosomes, or combinations of multiple errors were observed and scored. Criteria for scoring errors are described in the immunofluorescence section.

### Drug Treatments

For immunofluorescence, cells were treated with 0.1% DMSO, UMK57 or UMK95 (Dr. B Kwok, University of Montreal) at the concentrations specified for 45 min prior to fixation. For time-lapse live-cell imaging, hPSCs were cultured in phenol-free mTeSR1 supplemented with 0.1% DMSO, 3 µM, 6 µM or 20 µM proTAME (Tocris) or 2 µM UMK57 or UMK95 during imaging. RPE-1 H2B-GFP cells were cultured in phenol-free media supplemented with 0.1% DMSO, 3 µM or 6 µM proTAME (Tocris). For monastrol arrest and release experiments, RPE-1 H2B-GFP cells were arrested in 100 µM Monastol (Tocris) for 6 hrs followed by washout with phenol-free media into 0.1% DMSO, 3 µM or 6 µM proTAME (Tocris).

### All-*trans* Retinoic Acid Differentiation Assay

H1 and H9 hESCs were plated on Matrigel-coated 18mm glass coverslips and grown in mTeSR1 in 12-well cell culture plates. After 24 h,1 µM all-*trans* retinoic acid (RA)(Sigma #R2625) was added to fresh mTeSR1 media. hPSCs were treated with daily media changes including 1 µM RA for 4 days prior to fixation. Daily morphological changes were monitored by bright-field phase contrast microscopy using a Hamamatsu ORCA®-Fusion Gen III Scientific CMOS camera mounted on a Nikon Eclipse Ti2E microscope with a Nikon CFI Super Plan Fluor LWD 20× ADM, 0.7 numerical aperture air objective. Image series in the *Z*-axis were obtained using 1 µm optical sections. Image acquisition and analysis were performed using NIS Elements (Nikon) and ImageJ (NIH). Representative images shown in Supplementary Fig. 4 are from selected single *Z*-plane that best illustrates the morphology.

### Statistics

GraphPad Prism was used for all statistical analysis. Statistical details can be found in the figure legends which describe the statistical tests used and corresponding n values. Error bars represent standard deviation (SD). All experiments were performed in three or more replicates. Significance was defined as *p < 0.05, **p < 0.01, ***p<0.001, ****p < 0.0001. No outliers were excluded in data analysis.

## Acknowledgements

We thank Jonathan Draper, Aaron Straight, Bruce Conklin, Benjamin Kwok, and Thorsten Schlaeger for providing reagents or technical advice. We thank Ann Lavanway and the Dartmouth College Life Sciences Light Microscopy Facility for assistance with microscopy. Also, we thank members of the Compton and Godek laboratories for their helpful discussions and comments, particularly Thomas Kucharski for his suggestions. This work was supported by grant funding from the National Institutes of Health GM051542 to DAC and R01HD101436 to KMG and a Hitchcock Foundation Pilot grant to KMG.

## Author Contributions

Conceptualization-KMG; Methodology-KMG, CD, and AY; Validation-KMG, CD and AY; Formal Analysis-KMG, CD and AY; Investigation-KMG, CD and AY; Resources-KMG and DAC; Writing-Original Draft-KMG and CD; Writing-Review and Editing-KMG, CD, DAC and AY; Visualization-KMG and CD; Supervision-KMG; Funding Acquisition-KMG and DAC.

