## Supplemental Files for "A Pluripotent Developmental State Confers a Low Fidelity of Chromosome Segregation"

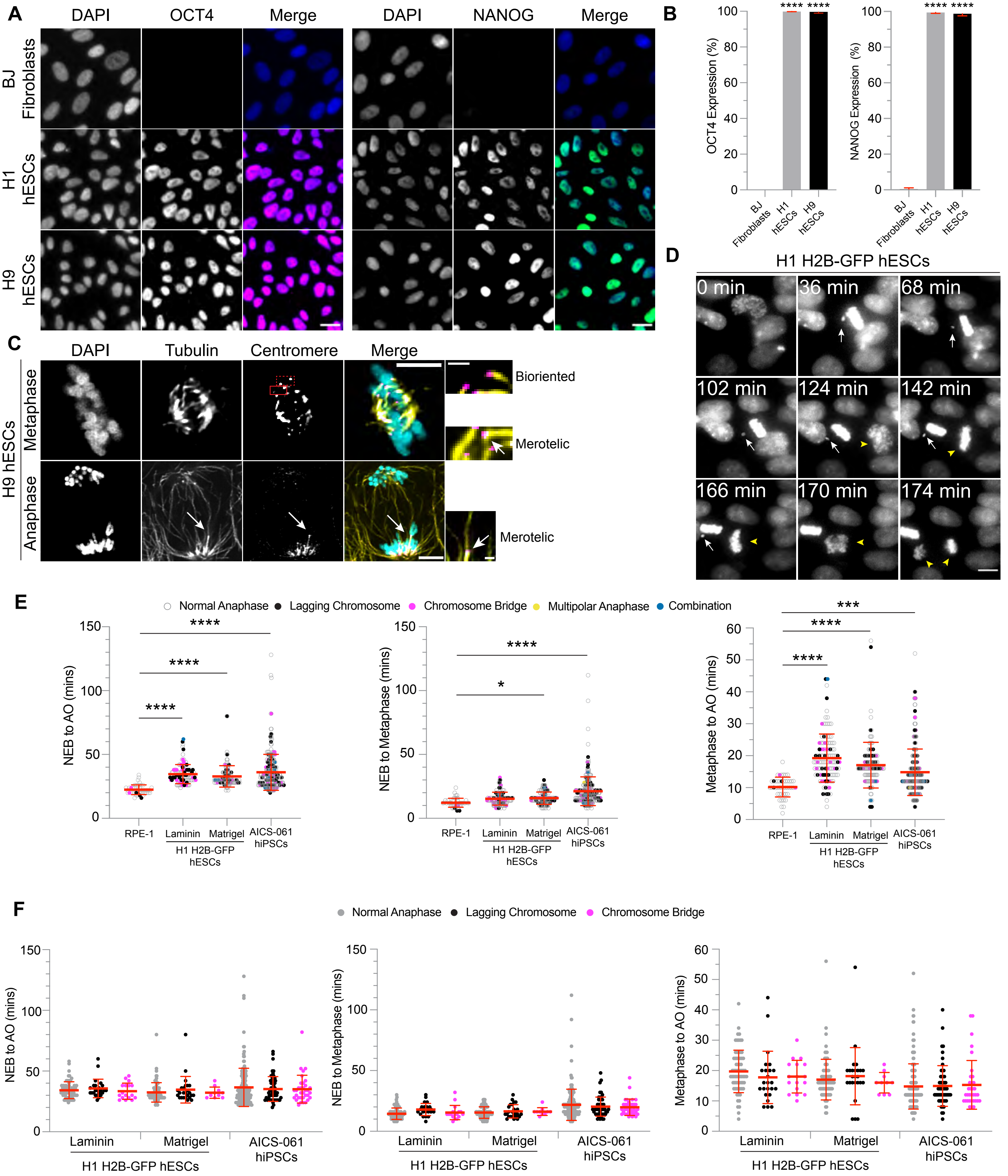
A
DAPI
OCT4
Merge
NANOG
BJ Fibroblasts
H1 hESCs
H9 hESCs
B
OCT4 Expression (%)
NANOG Expression (%)
**** ****
C
Tubulin
Centromere
H9 hESCs
Metaphase
Anaphase
Bioriented
Merotelic
D
H1 H2B-GFP hESCs
0 min
36 min
68 min
102 min
124 min
142 min
166 min
170 min
174 min
E
Normal Anaphase
Lagging Chromosome
Chromosome Bridge
Multipolar Anaphase
Combination
NEB to AO (mins)
NEB to Metaphase (mins)
Metaphase to AO (mins)
RPE-1
Laminin
Matrigel
AICS-061 hiPSCs
H1 H2B-GFP hESCs
F
Normal Anaphase
Lagging Chromosome
Chromosome Bridge

Figure S1

**Figure S1. Pluripotency transcription factor expression, SAC function and mitotic durations in hPSCs. Related to Figure 1.** (**A**) Representative images of primary somatic BJ fibroblasts, H1 and H9 hESCs that were stained with DAPI (blue) and OCT4 (magenta) or NANOG (green). Scale bars: 20 μm. (**B**) Percentage of cells expressing OCT4 or NANOG. n = 330 (BJ fibroblasts, OCT4), 1459 (H1, OCT4), 1103 (H9, OCT4), 389 (BJ fibroblasts, NANOG), 1183 (H1, NANOG), and 889 (H9, NANOG) cells from three independent experiments; mean ± SD, **** $p < 0.0001$ using a one-way ANOVA and Dunnett's multiple comparisons test. (**C**) Representative images of chromosome microtubule attachments in metaphase and anaphase H9 hESCs. Shown is DNA (cyan), microtubules (yellow) and centromeres (magenta). In the metaphase cell, boxes are pairs of centromeres with a bioriented attachment (dashed) or a merotelic attachment (solid, white arrow). The anaphase cell shows a lagging chromosome with a merotelic attachment (white arrow). Insets are magnified views. Scale bars: 5 μm (main) and 1 μm (insets). (**D**) Selected panels from time-lapse live-cell fluorescent imaging of H1 H2B-GFP hESCs showing a cell that maintained a mitotic arrest due to an unaligned chromosome (white arrows) and an adjacent cell that progressed through metaphase (yellow arrowheads). The unaligned chromosome is contrasted for better visualization. Scale bar: 10 μm. (**E**) Mitotic duration (left), prometaphase duration (middle) and metaphase duration (right) of somatic RPE-1 H2B-GFP, H1 H2B-GFP hESCs plated as single cells on Laminin-521 or as aggregates on Matrigel and AICS-061 hiPSCs. n = 46 anaphases in RPE-1 and n = 111 (H1 on Laminin-521), 121 (H1 on Matrigel), and 258 (AICS-061) anaphases in hPSCs from at least three independent experiments; NEB: nuclear envelope breakdown; AO: anaphase onset; mean ± SD; *$p < 0.05$, ***$p < 0.001$, **** $p < 0.0001$ using a one-way ANOVA and Dunnett's multiple comparisons test. (**F**) Mitotic duration, prometaphase and metaphase durations from panel E separated into groups of normal anaphases, anaphases with lagging chromosomes or anaphases with chromosome bridges; mean ± SD; a one-way ANOVA and Dunnett’s multiple comparisons test was used to test for significance.

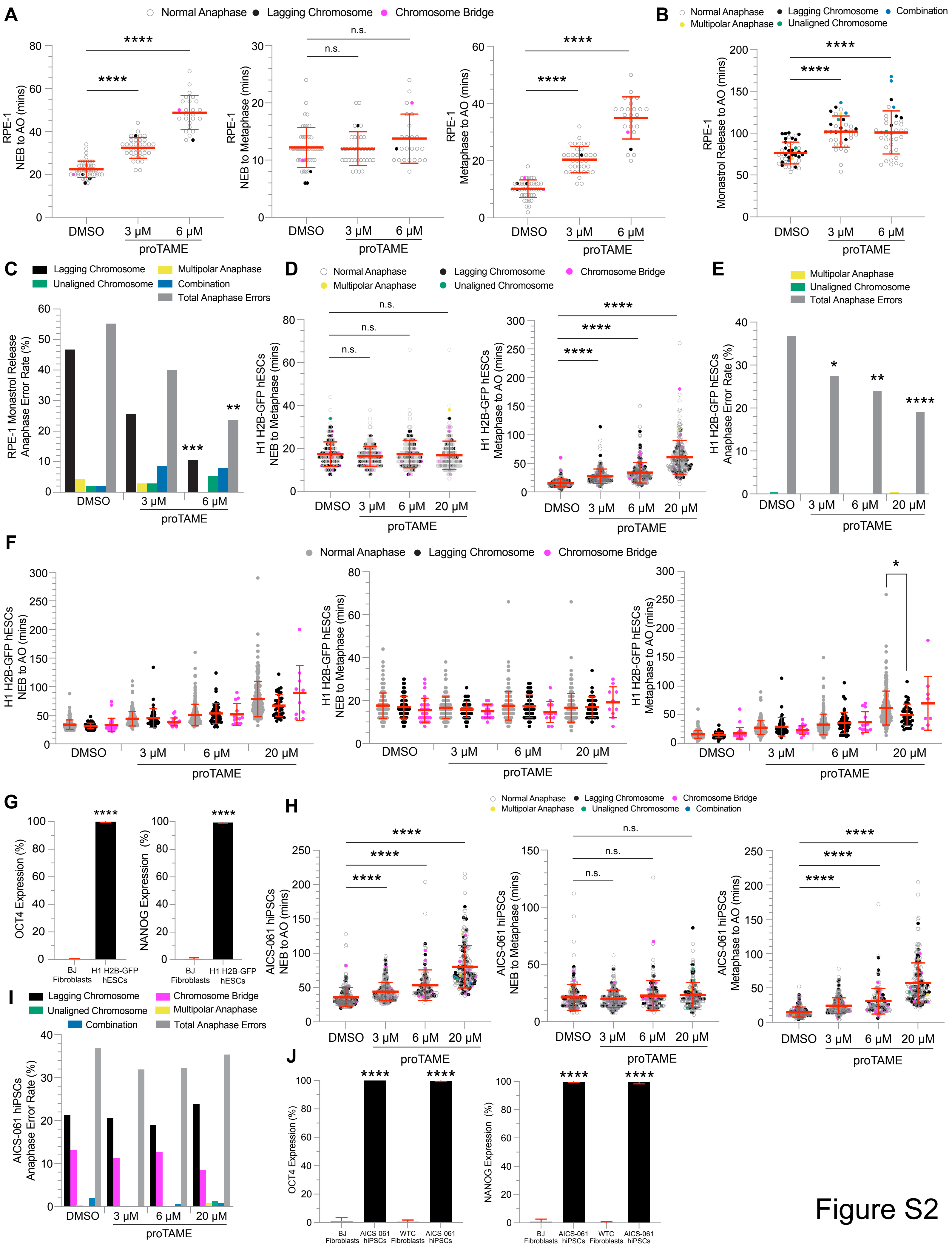
A
Normal Anaphase
Lagging Chromosome
Chromosome Bridge
RPE-1 NEB to AO (mins)
RPE-1 NEB to Metaphase (mins)
RPE-1 Metaphase to AO (mins)
DMSO
3 µM
6 µM
proTAME
n.s.
****
B
Normal Anaphase
Lagging Chromosome
Combination
Multipolar Anaphase
Unaligned Chromosome
RPE-1 Monastrol Release to AO (mins)
C
Lagging Chromosome
Multipolar Anaphase
Unaligned Chromosome
Combination
Total Anaphase Errors
RPE-1 Monastrol Release Anaphase Error Rate (%)
***
**
D
H1 H2B-GFP hESCs NEB to Metaphase (mins)
H1 H2B-GFP hESCs Metaphase to AO (mins)
20 µM
E
H1 H2B-GFP hESCs Anaphase Error Rate (%)
*
F
H1 H2B-GFP hESCs NEB to AO (mins)
G
OCT4 Expression (%)
NANOG Expression (%)
BJ Fibroblasts
H1 H2B-GFP hESCs
H
AICS-061 hiPSCs NEB to AO (mins)
AICS-061 hiPSCs NEB to Metaphase (mins)
AICS-061 hiPSCs Metaphase to AO (mins)
I
Chromosome Bridge
AICS-061 hiPSCs Anaphase Error Rate (%)
J
AICS-061 hiPSCs
WTC Fibroblasts
Figure S2

**Figure S2. Mitotic durations, anaphase error rates and pluripotency transcription factor expression in somatic cells or hPSCs. Related to Figure 2.** (**A**) Mitotic duration (left), prometaphase duration (middle) and metaphase duration (right) of somatic RPE-1 H2B-GFP cells treated with DMSO or increasing concentrations of proTAME. n = 46 (DMSO), 36 (3 μM proTAME), and 25 (6 μM proTAME) anaphases from three independent experiments; NEB: nuclear envelope breakdown; AO: anaphase onset; mean ± SD; n.s. $p > 0.05$, ****$p < 0.0001$ using a one-way ANOVA and Dunnett's multiple comparisons test. The data for DMSO control treated RPE-1 H2B-GFP cells is also shown in Figures 1E and S1E. (**B**) Time from monastol release to anaphase onset of somatic RPE-1 H2B-GFP cells treated with DMSO or increasing concentrations of proTAME. n = 47 (DMSO), 35 (3 μM proTAME), and 38 (6 μM proTAME) anaphases from three independent experiments; mean ± SD; **** $p < 0.0001$ using a one-way ANOVA and Dunnett's multiple comparisons test. (**C**) Percentage of anaphase errors in somatic RPE-1 H2B-GFP cells from panel B. n = 47 (DMSO), 35 (3 μM proTAME), and 38 (6 μM proTAME) anaphases from three independent experiments; **$p < 0.01$, ***$p < 0.001$ using a two-tailed Fisher's exact test. (**D**) Prometaphase duration (left) and metaphase duration (right) of H1 H2B-GFP hESCs treated with DMSO or increasing concentrations of proTAME from Figure 2C. n = 275 (DMSO), 247 (3 μM proTAME), 250 (6 μM proTAME), and 257 (20 μM proTAME) anaphases from six independent experiments; NEB: nuclear envelope breakdown; AO: anaphase onset; mean ± SD; n.s. $p > 0.05$, ****$p < 0.0001$ using a one-way ANOVA and Dunnett's multiple comparisons test. (**E**) Percentage of total anaphase errors, unaligned chromosomes, and multipolar anaphases in H1 H2B-GFP hESCs from Figure 2C. n = 275 (DMSO), 247 (3 μM proTAME), 250 (6 μM proTAME), and 257 (20 μM proTAME) anaphases from six independent experiments; *$p < 0.05$, **$p < 0.01$, ****$p < 0.0001$ using a two-tailed Fisher's exact test. (**F**) Mitotic duration (left), prometaphase (middle) and metaphase (right) durations from Figure 2C and panel D separated into groups of normal anaphases, anaphases with lagging chromosomes or anaphases with chromosome bridges. n = 275 (DMSO), 247 (3 μM proTAME), 250 (6 μM proTAME), and 257 (20 μM proTAME) anaphases from six independent experiments; NEB: nuclear envelope breakdown; AO: anaphase onset; mean ± SD; * $p < 0.05$ using a one-way ANOVA and Dunnett's multiple comparisons test. (**G**) Percentage of cells expressing OCT4 or NANOG in parallel samples from Figure 2C. n = 600 cells from six independent experiments; mean ± SD; **** $p < 0.0001$ using a two-tailed Student's t test. (**H**) Mitotic duration (left), prometaphase duration (middle) and metaphase duration (right) of AICS-061 hiPSCs treated with DMSO or increasing concentrations of proTAME. n = 258 (DMSO), 282 (3 μM proTAME), 158 (6 μM proTAME), and 226 (20 μM proTAME) anaphases from six independent experiments; NEB: nuclear envelope breakdown; AO: anaphase onset; mean ± SD; n.s. $p > 0.05$, ****$p < 0.0001$ using a one-way ANOVA and Dunnett's multiple comparisons test. The data for DMSO control treated AICS-061 hiPSCs is also shown in Figures 1E, S1E-F. (**I**) Percentage of anaphase errors in AICS-061 hiPSCs treated with DMSO or increasing concentrations of proTAME from panel H. n = 258 (DMSO), 282 (3 μM proTAME), 158 (6 μM proTAME), and 226 (20 μM proTAME) anaphases from six independent experiments; two-tailed Fisher's exact test was used to test for significance. The data for DMSO control treated AICS-061 hiPSCs is also shown in Figure 1E. (**J**) Percentage of cells expressing OCT4 or NANOG in parallel samples from panel H. n = 300 cells from six independent experiments with the first three replicates comparing AICS-061 hiPSCs to BJ fibroblasts and the last three replicates comparing AICS-061 hiPSCs to isogenic WTC-11 fibroblasts; mean ± SD; **** $p < 0.0001$ using two-tailed Student's t test.

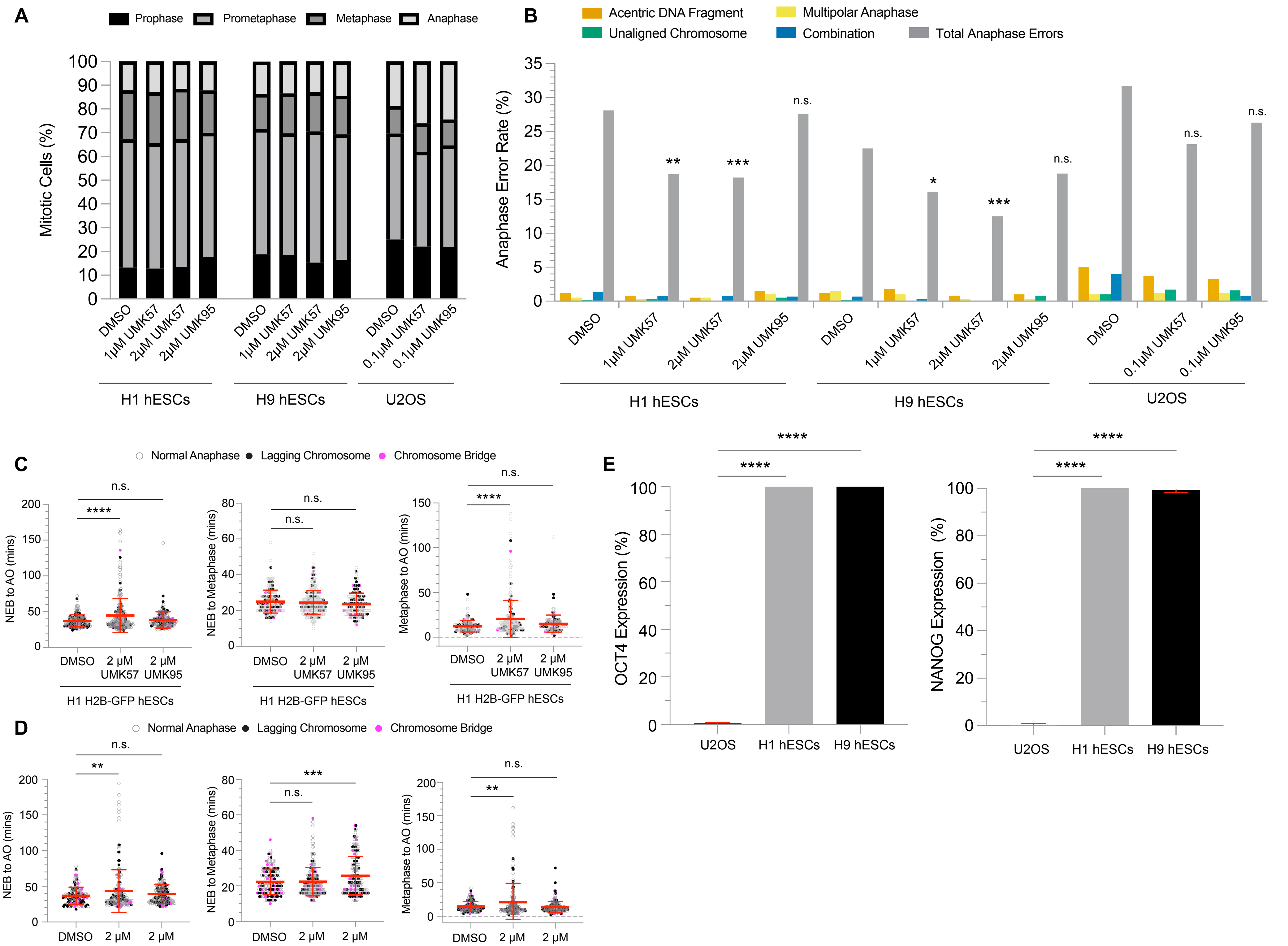

A
Prophase
Prometaphase
Metaphase
Anaphase
Mitotic Cells (%)
DMSO
1µM UMK57
2µM UMK57
2µM UMK95
0.1µM UMK57
0.1µM UMK95
H1 hESCs
H9 hESCs
U2OS
B
Acentric DNA Fragment
Unaligned Chromosome
Multipolar Anaphase
Combination
Total Anaphase Errors
Anaphase Error Rate (%)
n.s.
**
***
*
C
Normal Anaphase
Lagging Chromosome
Chromosome Bridge
NEB to AO (mins)
NEB to Metaphase (mins)
Metaphase to AO (mins)
****
2 µM UMK57
2 µM UMK95
H1 H2B-GFP hESCs
D
AICS-061 hiPSCs
E
OCT4 Expression (%)
NANOG Expression (%)
U2OS


Figure S3

**Figure S3. Mitotic progression, anaphase error rates, mitotic durations and pluripotency transcription factor expression in U2OS cancer cells or hPSCs. Related to Figure 3.** (**A**) Percentage of mitotic cells in prophase, prometaphase, metaphase or anaphase in H1 and H9 hESCs and U2OS cancer cells treated with DMSO, UMK57 or UMK95. n = 3562 (H1, DMSO), 3010 (H1, 1 μM UMK57), 3389 (H1, 2 μM UMK57), 3360 (H1, 2 μM UMK95), 2949 (H9, DMSO), 2869 (H9, 1 μM UMK57), 2830 (H9, 2 μM UMK57), 2713 (H9, 2 μM UMK95) cells in hESCs and n = 1059 (DMSO), 930 (0.1 μM UMK57), and 993 (0.1 μM UMK95) cells in U2OS. There was no significant difference in the proportion of prometaphase cells for hESCs treated with DMSO, UMK57 or UMK95; two-tailed Fisher's exact test. (**B**) Percentage of acentric DNA fragments, unaligned chromosomes, multipolar anaphases and total anaphase errors in H1 and H9 hESCs and U2OS cancer cells from Figure 3B treated with DMSO, UMK57 or UMK95. n = 431 (DMSO), 390 (1 μM UMK57), 391 (2 μM UMK57), and 410 (2 μM UMK95) anaphases in H1 hESCs. n = 405 (DMSO), 385 (1 μM UMK57), 367 (2 μM UMK57), and 393 (2 μM UMK95) anaphases in H9 hESCs. n = 199 (DMSO), 242 (0.1 μM UMK57), and 243 (0.1 μM UMK95) anaphases in U2OS from three independent experiments; n.s. $p > 0.05$, *$p < 0.05$, **$p < 0.01$, ***$p < 0.001$ using a two-tailed Fisher's exact test. (**C**) Mitotic duration (left), prometaphase (middle) and metaphase (right) durations from time-lapse live-cell fluorescent imaging of H1 H2B-GFP hESCs in Figure 3C treated with DMSO, UMK57 or UMK95. n = 205 (DMSO), 278 (2 μM UMK57), and 187 (2 μM UMK95) anaphases from three independent experiments; NEB: nuclear envelope breakdown; AO: anaphase onset; n.s. $p > 0.05$, ****$p < 0.0001$ using a one-way ANOVA and Dunnett's multiple comparisons test. (**D**) Mitotic duration (left), prometaphase (middle) and metaphase (right) durations from time-lapse live-cell fluorescent imaging of AICS-061 hiPSCs in Figure 3D treated with DMSO, UMK57 or UMK95. n = 158 (DMSO), 209 (2 μM UMK57), and 177 (2 μM UMK95) anaphases from three independent experiments; NEB: nuclear envelope breakdown; AO: anaphase onset; n.s. $p > 0.05$, **$p< 0.01$, ***$p < 0.001$ using a one-way ANOVA and Dunnett's multiple comparisons test. (**E**) Percentage of cells expressing OCT4 or NANOG in parallel samples from Figure 3B. $n = 300$ cells from three independent experiments; mean ± SD; **** $p < 0.0001$ using a one-way ANOVA and Dunnett's multiple comparisons test.

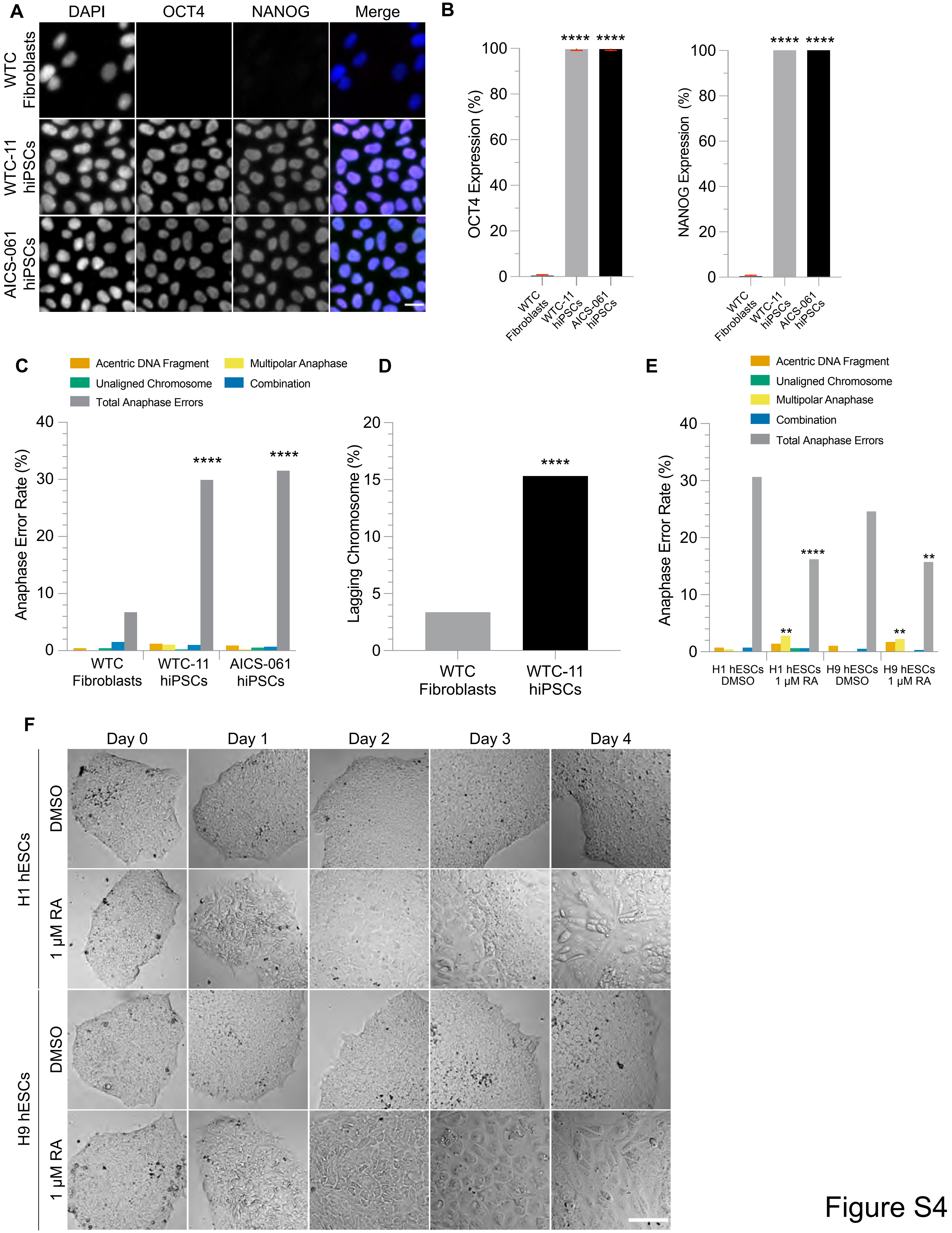
A
DAPI
OCT4
NANOG
Merge
WTC Fibroblasts
WTC-11 hiPSCs
AICS-061 hiPSCs
B
OCT4 Expression (%)
NANOG Expression (%)
****
WTC Fibroblasts
WTC-11 hiPSCs
AICS-061 hiPSCs
C
Acentric DNA Fragment
Multipolar Anaphase
Unaligned Chromosome
Combination
Total Anaphase Errors
Anaphase Error Rate (%)
D
Lagging Chromosome (%)
E
H1 hESCs DMSO
H1 hESCs 1 µM RA
H9 hESCs DMSO
H9 hESCs 1 µM RA
F
Day 0
Day 1
Day 2
Day 3
Day 4
H1 hESCs
H9 hESCs
DMSO
1 µM RA

Figure S4

**Figure S4. Pluripotency transcription factor expression, anaphase error rates and morphology during RA treatment in somatic cells or hPSCs. Related to Figure 4.** (**A**) Representative images of isogenic WTC fibroblasts, WTC-11 hiPSCs and AICS-061 hiPSCs that were stained with DAPI (blue), OCT4 (magenta) and NANOG (green). Scale bar: 20 μm. (**B**) Percentage of cells expressing OCT4 or NANOG. n = 300 cells from three independent experiments; mean ± SD; **** $p < 0.0001$ using a one-way ANOVA and Dunnett's multiple comparisons test. (**C**) Percentage of acentric DNA fragments, unaligned chromosomes, multipolar anaphases and total anaphase errors in isogenic WTC fibroblasts, WTC-11 hiPSCs and AICS-061 hiPSCs from Figure 4B. n = 268 (WTC fibroblasts), 421 (WTC-11 hiPSCs), and 438 (AICS-061 hiPSCs) anaphases from three independent experiments; ****$p < 0.0001$ using a two-tailed Fisher's exact test. (**D**) The lagging chromosome rate in WTC-11 hiPSCs remains significantly increased compared to isogenic WTC-11 fibroblasts after hypothetically excluding 10% of the lagging chromosome data due to aneuploid WTC-11 hiPSCs; ****$p < 0.0001$ using a two-tailed Fisher's exact test. (**E**) Percentage of acentric DNA fragments, unaligned chromosomes, multipolar anaphases and total anaphase errors in H1 and H9 hESCs from Figure 4C after treatment with DMSO or 1 μM all-*trans* retinoic acid (RA) for 4 days. n = 454 (H1, DMSO), 358 (H1, 1 μM RA), 398 (H9, DMSO), and 356 (H9, 1 μM RA) anaphases from three independent experiments; **$p < 0.01$, ****$p < 0.0001$ using a two-tailed Fisher's exact test. (**F**) Representative phase contrast images of daily morphological changes during RA treatment in H1 and H9 hESCs. Scale bar: 100 μm.

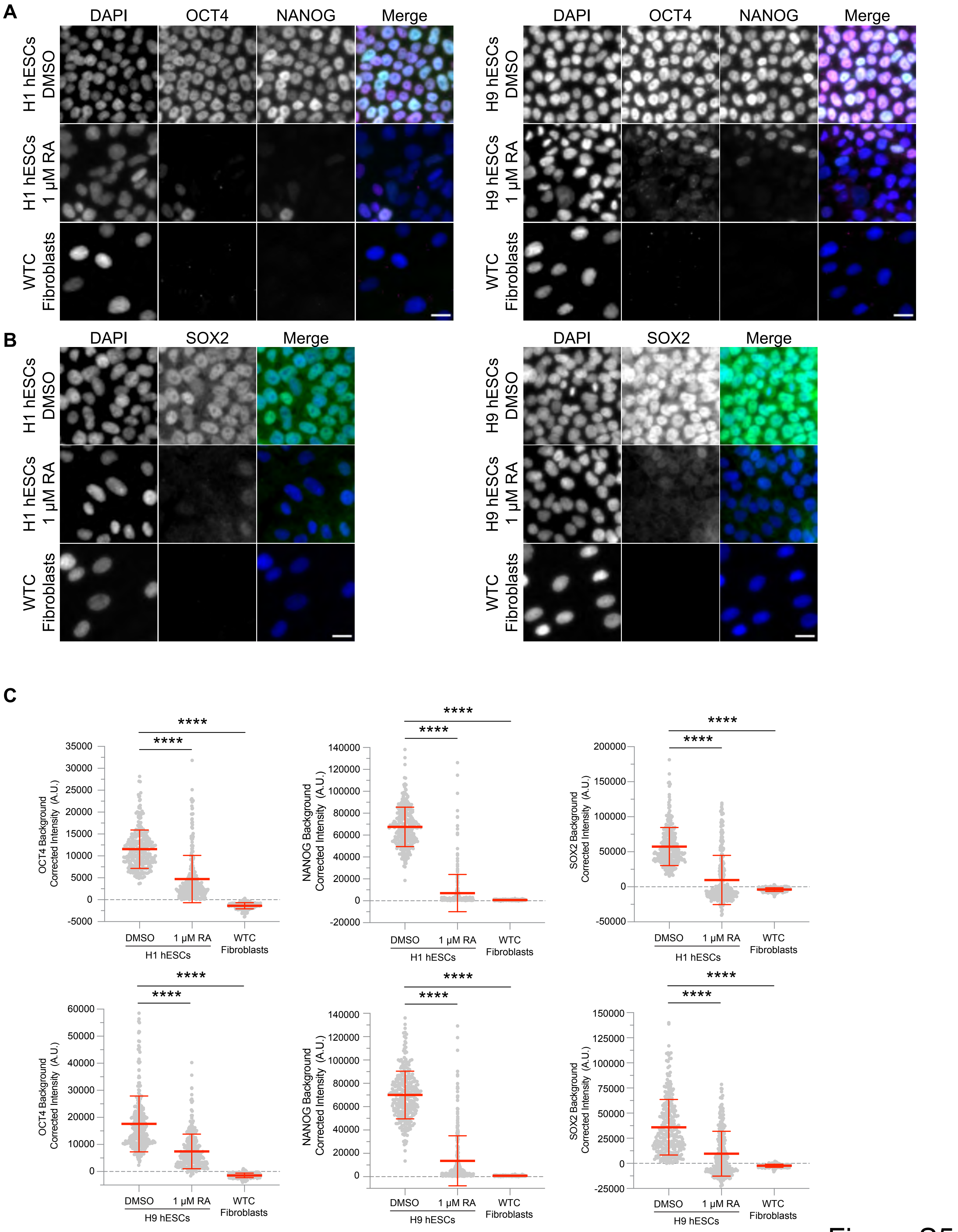
A
DAPI
OCT4
NANOG
Merge
H1 hESCs DMSO
H1 hESCs 1 µM RA
WTC Fibroblasts
H9 hESCs DMSO
H9 hESCs 1 µM RA
B
SOX2
C
****
OCT4 Background Corrected Intensity (A.U.)
NANOG Background Corrected Intensity (A.U.)
SOX2 Background Corrected Intensity (A.U.)
DMSO
1 µM RA
WTC Fibroblasts
H1 hESCs
H9 hESCs

Figure S5

**Figure S5. Quantification of OCT4, NANOG and SOX2 protein levels in H1 and H9 hESCs upon RA treatment. Related to Figure 4.** (**A**) Representative images of H1 and H9 hESCs treated with DMSO or 1 μM all-*trans* retinoic acid (RA) for 4 days and WTC fibroblasts that were stained with DAPI (blue), OCT4 (magenta) and NANOG (green). Scale bars: 20 μm. (**B**) Representative images of H1 and H9 hESCs treated with DMSO or 1 μM RA for 4 days and WTC fibroblasts that were stained with DAPI (blue) and SOX2 (green). Scale bars: 20 μm. (**C**) Quantification of OCT4, NANOG and SOX2 protein levels in H1 and H9 hESCs and WTC fibroblasts. $n = 300$ cells from three independent experiments; mean ± SD; **** $p < 0.0001$ using a one-way ANOVA and Dunnett's multiple comparisons test.

**Video S1. Movie of the H1 H2B-GFP hESC showing a normal anaphase. Related to Figure 1D.**

**Video S2. Movie of the H1 H2B-GFP hESC showing an erroneous anaphase with a lagging chromosome. Related to Figure 1D.**

**Video S3. Movie of the H1 H2B-GFP hESC showing an erroneous anaphase with a chromosome bridge. Related to Figure 1D.**

**Video S4. Movie of the H1 H2B-GFP hESC maintaining a mitotic arrest due to an unaligned chromosome and an adjacent cell progressing through metaphase. Related to Figure S1D.**
